# GPLD1 Regulates the Shedding of JUNO to Block Polyspermy in Porcine Oocyte

**DOI:** 10.64898/2026.04.01.706704

**Authors:** Boda Chen, Luting Shi, Fan Xia, Xingguo Chen, Jingyi Wang, Sitong Gao, Xiaoqian Zhou, Pengyun Ji, Guoshi Liu, Lu Zhang

## Abstract

Mammalian fertilization commences with the essential interaction between sperm IZUMO1 and its oocyte-surface receptor, JUNO. Following gamete fusion, JUNO is rapidly shed from the oocyte to establish a definitive membrane block to polyspermy, a pathological condition that remains a major hurdle in porcine *in vitro* fertilization (IVF). Despite its biological importance, the molecular networks driving JUNO cleavage has remained elusive. Here, by integrating proteomics, dual-color live-cell imaging, and functional perturbations, we identify the extended existence of JUNO and the GPI-specific phospholipase D1 (GPLD1) as the requisite enzyme mediating JUNO shedding in porcine oocytes. Targeted knockdown or pharmacological inhibition of GPLD1 stabilizes oocyte JUNO, prolongs the window of oocyte receptivity, and significantly exacerbates polyspermy, ultimately compromising embryonic developmental competence. Conversely, GPLD1 overexpression restricts redundant sperm adherence and enhances the efficiency of monospermic zygote formation and blastocyst development. Live-cell imaging reveals that fertilization triggers a transient, pulsed recruitment of GPLD1 in the oocyte, which precisely coincides with the biphasic kinetics of JUNO depletion. Our findings establish that the enzymatic cleavage of the GPI-anchor by GPLD1 is critical for JUNO release, defining a fundamental mechanism for the membrane-level block to polyspermy. This work provides a molecular framework for ensuring sperm-oocyte recognition and improving in vitro fertilization outcomes in mammals.

## Introduction

Fertilization is a highly orchestrated cascade of molecular and cellular events culminating in the recognition, adhesion, and fusion of a single spermatozoon with an oocyte to form a zygote. Prior to membrane fusion, sperm must complete capacitation and the acrosome reaction to penetrate the zona pellucida (ZP)-the specialized glycoprotein matrix surrounding the oocyte (Bleil and Wassarman., 1980; Prasad et al., 2000; Boja et al., 2003). The molecular and structural basis of sperm–ZP recognition and removement post fertilization have been extensively characterized, with specific ZP glycoproteins and ovastacin identified as essential mediators (Burkart et al., 2012; Nishio et al., 2024). Far less is understood about how sperm–oocyte interactions are regulated at the plasma membrane, where final binding and fusion occur. Current insights into membrane-level gamete recognition derive primarily from murine and human models, leaving the regulatory mechanisms operating in other mammalian oocytes comparatively unexplored (Inoue et al., 2003; Lorenzetti et al., 2014; Barbaux et al., 2020; Lamas-Toranzo et al., 2020; Fujihara et al., 2020, 2021; Noda et al., 2020, 2022; Matsumura et al., 2022; Tang et al., 2022).

The proteins located on membranes form the secondary species-specific molecular networks for recognition and fusion of sperm and oocyte. The identification of sperm IZUMO1 and its receptor in the oocyte membrane JUNO (also known as IZUMO1R) (Inoue et al., 2005; Bianchi et al., 2014), help us to elucidate the molecular mechanisms for this critical process. JUNO is a glycosylphosphatidylinositol (GPI)-anchored protein, and genetic ablation of Juno in mice abolishes sperm binding and fusion, resulting in complete female infertility (Bianchi et al., 2014). The IZUMO1–JUNO interaction is evolutionarily conserved and mediates species-specific gamete adhesion in mammals, including humans, mouse, porcine and bovine (Ohto et al., 2016; Kato et al., 2016; Aydin et al., 2016; Boult et al., 2025). Mutations in JUNO is related to ART failure due to polyspermy (Yu et al., 2018). Importantly, JUNO is rapidly shed from the oocyte into the perivitelline space shortly after fertilization, a process proposed to constitute a membrane-level block to polyspermy (Bianchi et al., 2014). However, despite its central importance, the molecular mechanism responsible for JUNO removal from the oocyte membrane remains unknown.

JUNO is a glycosylphosphatidylinositol-anchored protein (GPI-AP) and a member of the folate receptor family; however, it is functionally distinct, involved in molecular adhesion rather than acting as a transporter for folate (Chen et al., 2013; Wibowo et al., 2013). GPI-anchored proteins are commonly regulated through enzymatic cleavage of their GPI anchors, a process mediated by GPI-specific phospholipases (Udenfriend et al., 1995). Among these, GPI-specific phospholipase D1 (GPLD1) is a well-characterized mammalian enzyme capable of cleaving GPI anchors between inositol and phosphate, thereby releasing GPI-anchored proteins from the plasma membrane (Brown and Rose, 1992; Ikezawa, 2002). GPLD1 has been implicated in diverse biological processes, including membrane protein remodeling and signal transduction, yet its role in mammalian fertilization has not been examined (Horowitz et al., 2020; Ren et al., 2024). Notably, proteome studies indicate that GPLD1 is highly expressed in mammalian oocytes, particularly in oocytes derived from large follicles, and remains detectable throughout early embryogenesis (Wang et al., 2010; Li et al., 2022). These observations raise the intriguing possibility that GPLD1 may participate in the post-fertilization remodeling of the oocyte by regulating the shedding of GPI-anchored proteins such as JUNO.

Successful embryonic development requires establishing a robust block to polyspermy, as the entry of multiple sperm invariably causes chromosomal imbalance and early embryonic arrest (Nguyen et al., 2020). Although polyspermy is generally rare in most mammals, porcine fertilization presents a striking exception (Suzuki et al., 2003; Nagai and Moor, 1990). In pigs, polyspermy occurs in 30–40% of *in vivo* fertilizations even exceed 75% under *in vitro* conditions (Romar et al., 2016; Mahe et al., 2021), representing a major barrier to reproductive efficiency for sow. Despite its biological and agricultural significance, the molecular mechanisms underlying the exceptional susceptibility of porcine oocytes to polyspermy remain largely unknown.

Here, we address this hypothesis by combining ultra-sensitive proteomic profiling with high-resolution imaging and functional perturbation approaches to dissect membrane protein dynamics during porcine fertilization. We identify JUNO and GPLD1 as key components of a membrane regulatory axis that governs sperm–egg recognition, polyspermy prevention, and early embryonic development. By elucidating the spatiotemporal dynamics and functional interplay of JUNO and GPLD1, our study reveals a previously unrecognized mechanism of oocyte remodeling that contributes to fertilization fidelity and provides mechanistic insight into the unusually high incidence of polyspermy in pigs.

## Results

### The Expression and Localization of JUNO in Mammalian Oocytes

To investigate the expression patterns of the JUNO gene in oocytes and early embryos across different species, we utilized the GametesOmics (Yan et al., 2021) and DevOmics (An et al., 2024) databases, along with the raw transcriptomic data (Zhi et al., 2022). As shown in Fig. 1A-C, *JUNO* gene expression was detected in oocytes and early embryos of humans, mice, and porcine. In humans, compared to the fully-grown oocyte (FGO) stage, *JUNO* expression was lower at the growing oocyte (GO) stage, and remained expressed from fertilization through the 8-cell stage (Fig. 1 A). Similarly, porcine *JUNO* was expressed in oocytes and embryos up to the 8-cell stage (Fig. 1 B). In contrast, *Juno* showed high expression from the oocyte to zygote stages in mice, but rapidly declined after the 2-cell stage (Fig. 1 C). The JUNO expression in oocytes and early embryos is conserved across different species, but distinct differences in the timing of JUNO expression were observed among species, indicating that the function of JUNO requires further investigation.

**Figure 1.**
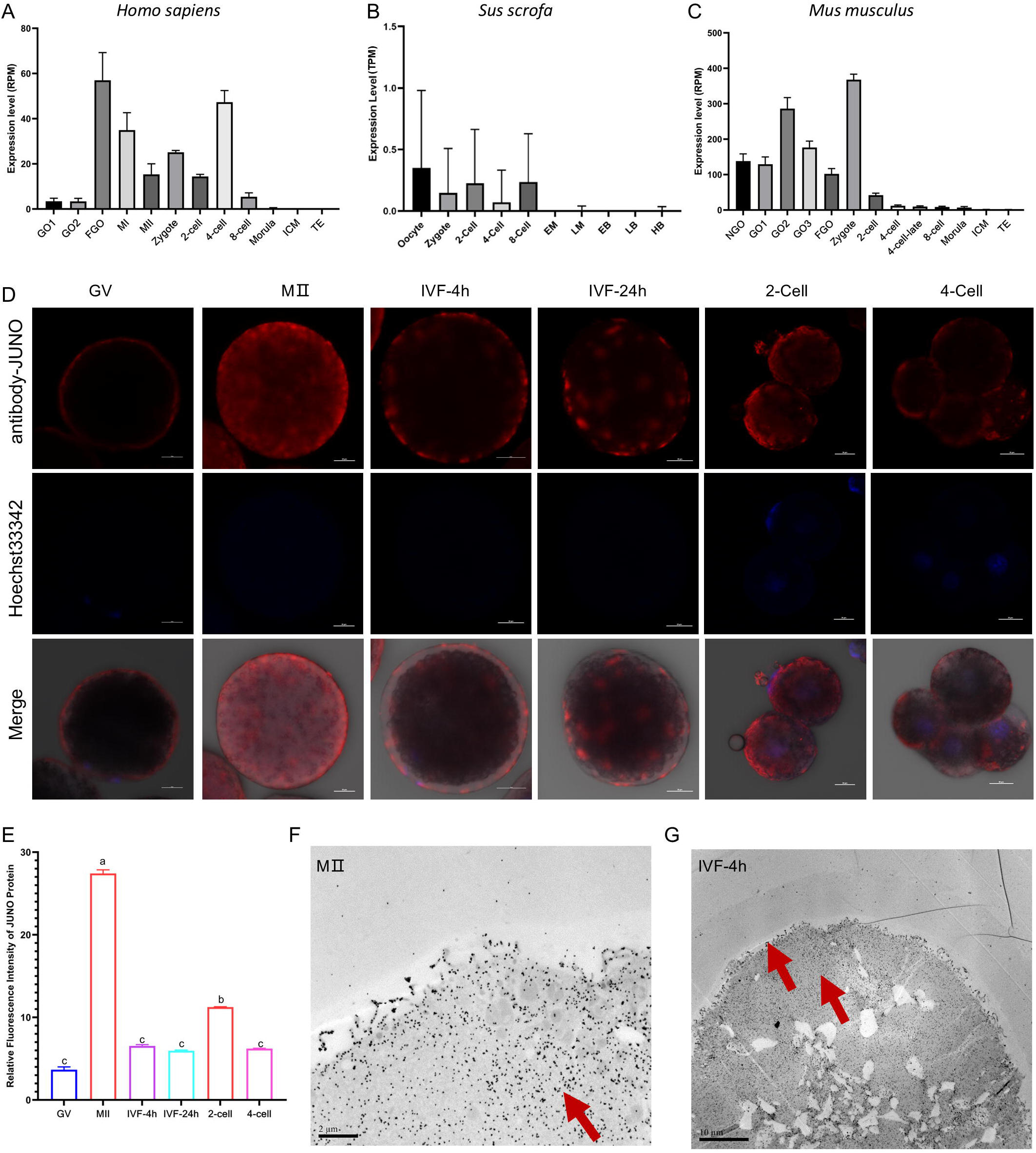
Spatiotemporal Expression Pattern of JUNO During Oocyte Maturation and Early Embryo Development. **(A-C)** Expression levels of JUNO across various developmental stages in *Homo sapiens* (A), *Mus musculus* (B), and *Sus scrofa* (C). **(D)** Representative immunofluorescence (IF) images showing the spatial distribution and abundance of JUNO (red) and DNA (Hoechst, blue) in porcine oocytes at the Germinal Vesicle (GV), MⅡ, IVF-4h, IVF-24h, 2-cell, and 4-cell stages. Scale bars, 20 μm. **(E)** Quantification of relative JUNO fluorescence intensity corresponding to the stages shown in (D). **(F and G)** Representative transmission electron microscopy (TEM) images showing the ultrastructural localization of JUNO on the membrane surface particles in porcine oocytes at the MⅡ and IVF-4h stages (red arrows indicate gold particles). Data are presented as mean ± SEM. Scale bars: 2 μm (MⅡ) and 10 μm (IVF-4h).

Then, the expression and localization of JUNO in porcine oocyte and early embryos were examined by immunofluorescence (IF) and transmission electron microscopy (TEM) (Fig. 1 D-G). It was confirmed that JUNO was predominantly localized on the oocyte and early embryonic membranes, displaying a ring-like distribution at the GV stage, and its expression is elevated at the MⅡ stage (Fig. 1 D-F). Post-fertilization, the JUNO drops significantly but remains at certain level till 4-cell stage (Fig. 1 D-F). Further analyses by TEM revealed that, relative to MⅡ oocytes, JUNO underwent a gradual redistribution toward the plasma membrane by 4 h post-fertilization (Fig. 1 G). Based on these observations, the persistent JUNO expression in porcine oocyte indicates that the shedding of JUNO is delayed or incomplete, thereby permit continued sperm recognition.

To determine whether these species-specific expression dynamics are mirrored at the primary sequence level, we performed a comparative analysis of JUNO across diverse mammals. Previous studies have confirmed that JUNO is an essential receptor for mammalian fertilization; however, our analysis highlights sequence divergence among species (Fig. S1 B). The results demonstrate that JUNO sequences exhibit heterogeneity, with pig and mouse JUNO sharing 64.61% sequence identity (Fig. S1 C). Despite these widespread amino acid substitutions, we identified two conserved N-glycosylation sites and a C-terminal GPI-anchor signal sequence. These conserved motifs likely represent the minimal structural requirements retained across mammalian evolution to ensure basic membrane anchoring and stability, while the surrounding variable regions may dictate species-specific fertilization mechanics.

### JUNO is a critical determinant of porcine sperm-oocyte recognition and fusion

To investigate the functional role of JUNO in porcine oocytes, an siRNA-mediated knockdown approach was employed during *in vitro* maturation. Microinjection of JUNO-targeting siRNAs (si-JUNO) or scrambled RNAs (Control) at 32 hours of maturation, JUNO protein levels on the oocyte significantly downregulated, as confirmed by immunofluorescence (Fig. 2 A and B). At 4 hours post-insemination (IVF-4h), JUNO-depleted oocytes exhibited a marked reduction in sperm adhesion compared to scrambled siRNA controls (Fig. 2 C and E). Furthermore, analysis at 12 hours post-insemination (IVF-12h) revealed that JUNO knockdown resulted in a significantly higher proportion of normal zygotes (2PN), accompanied by a decrease in percentage of polyspermy formation (Fig. 2 D and F). Furthermore, the oocytes show high developmental competence as the cleavage and blastocyst rate were all elevated (Fig. 3).

**Figure 2.**
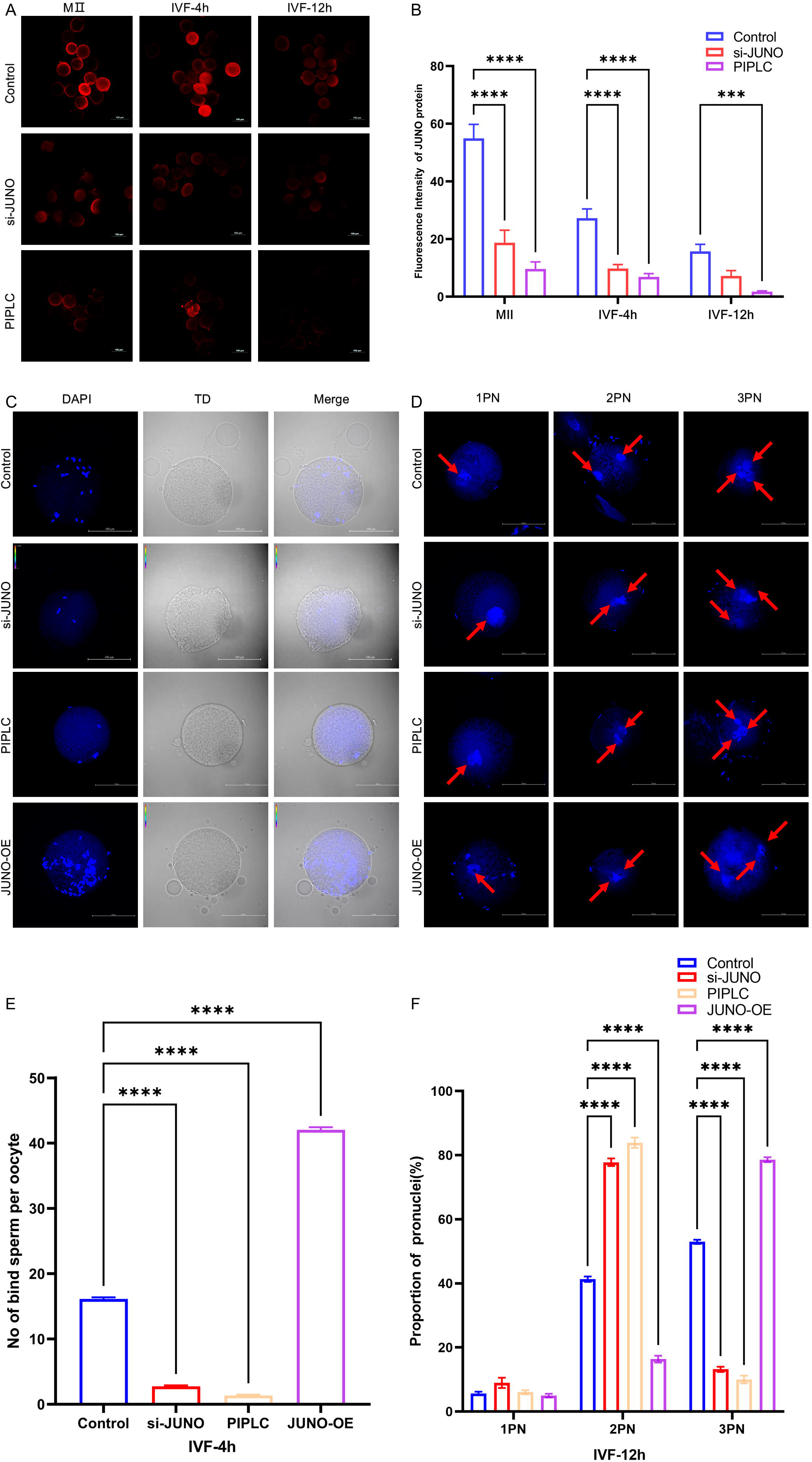
JUNO is Crucial for porcine Sperm Binding and Polyspermy Block. **(A)** Immunofluorescence images of JUNO in control group, PIPLC-treated, and si-JUNO oocytes at MⅡ, IVF-4h, and IVF-12h stages. **(B)** Quantitative analysis of JUNO fluorescence intensity showing the efficiency of JUNO depletion in the treatment groups. Data are presented as mean ± SEM; ***P < 0.001, ****P < 0.0001. Scale bars, 100 μm. **(C)** Representative fluorescence images showing sperm binding to porcine oocytes under different treatments (Control, si-JUNO, PIPLC, and JUNO-OE). Oocytes were stained with DAPI (blue) to visualize sperm nuclei. Scale bars, 100 μm. **(D)** Representative images of pronuclear formation in fertilized oocytes at 12 h post-IVF under different treatments. Arrows indicate pronuclei. Oocytes exhibiting one pronucleus (1PN), two pronuclei (2PN), or three pronuclei (3PN) were observed, reflecting variations in fertilization and polyspermy among the groups. Scale bars, 100 μm. **(E)** Bar chart comparing the number of bound sperm per oocyte at IVF-4h across Control, si-JUNO, PIPLC, and JUNO-OE (overexpression) groups. Data are presented as mean ± SEM.****P < 0.0001.**(F)** Proportion of pronuclei (1PN, 2PN, and polyspermic 3PN) observed at IVF-12h across the different experimental groups. Data are presented as mean ± SEM. ****P < 0.0001.

**Figure 3.**
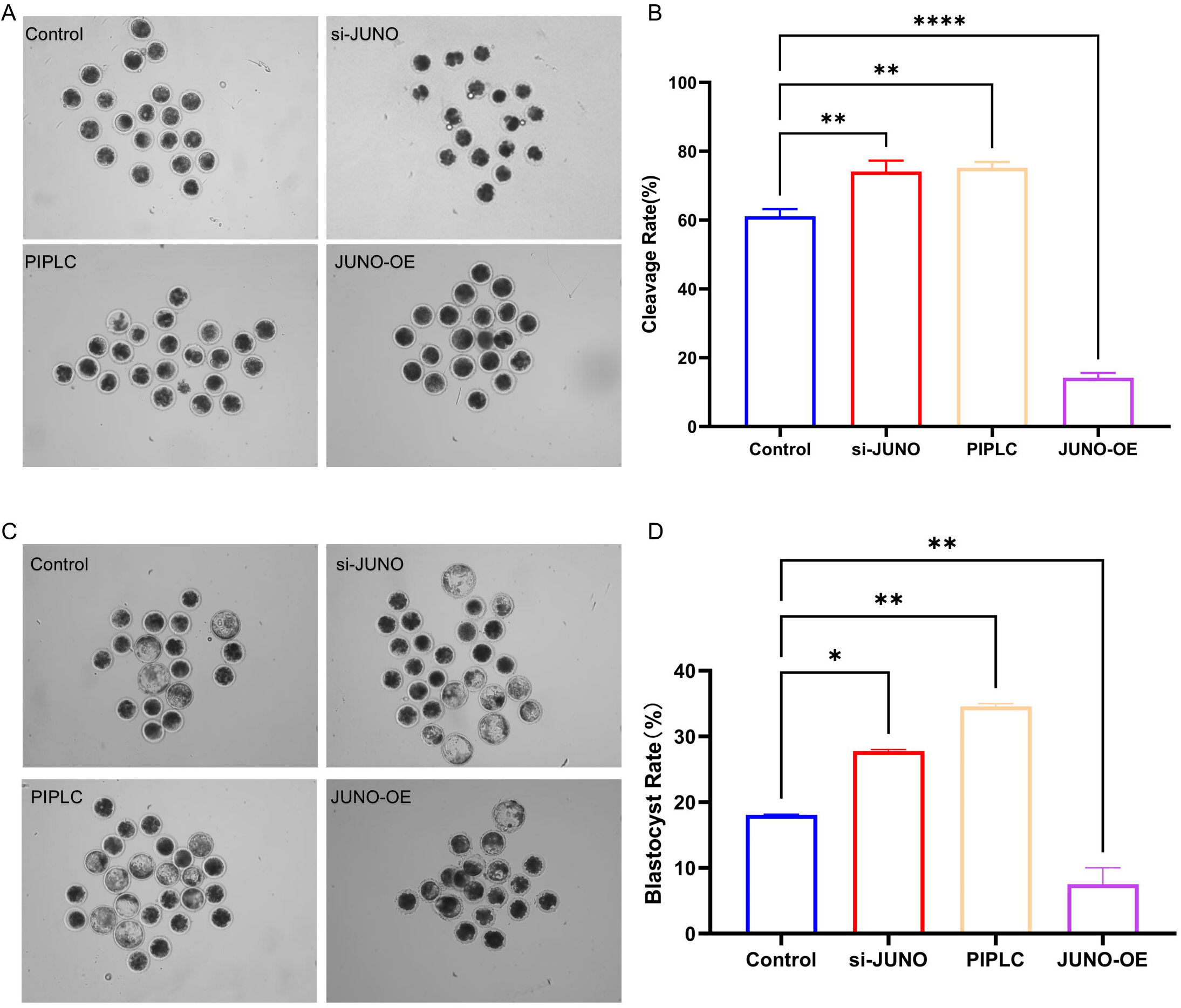
Effect of JUNO expression levels on early embryonic development. **(A)** Representative images of embryos at the cleavage stage. Micrographs showing the morphology of embryos in different treatment groups (Control, si-JUNO, PIPLC, and JUNO-OE) during early cleavage. **(B)** Comparison of cleavage rates across groups. Statistical analysis of the percentage of embryos reaching the cleavage stage. Data are presented as mean ± SEM.**P < 0.01, ****P < 0.0001. **(C)** Representative images of embryos at the blastocyst stage. Micrographs illustrating the development of embryos into blastocysts in the four experimental groups. **(D)** Comparison of blastocyst rates across groups. Statistical analysis of the percentage of embryos reaching the blastocyst stage. Data are presented as mean ± SEM. *P < 0.05, **P < 0.01.

Next, as JUNO is a glycosylphosphatidylinositol (GPI)-anchored protein, phosphatidylinositol-specific phospholipase C (PIPLC) was used to enzymatically disrupting its function in murine oocyte (Bianchi et al., 2014). Thus, zona-free porcine oocytes were treated with PIPLC, which specifically cleaves GPI anchors and releases associated proteins from the plasma membrane. Consistent with genetic knockdown data, PIPLC treatment resulted in a pronounced loss of JUNO signal from porcine oocytes (Fig. 2 A and B). This enzymatic removal of GPI-anchored proteins significantly diminished sperm binding capacity at 4 hours post-fertilization (Fig. 2 C and E). Moreover, frequency of 2PN zygotes formation at 12 hours increased and a corresponding reduction in rate of polyspermy (Fig. 2 D and F). Furthermore, the oocytes treated with PIPLC show high developmental competence as the cleavage and blastocyst rate were all elevated (Fig. 3).

Finally, we examined whether elevation of JUNO expression in porcine oocyte is sufficient to enhance sperm recruitment and alter fertilization outcomes. Following microinjection, oocytes overexpressing *JUNO* exhibited a dramatic elevation in sperm binding at 4 hours post-IVF compared to that in oocytes from control group (Fig. 2 C and E). However, JUNO-overexpressing zygotes showed a significant reduction in normal 2PN formation at 12 hours post-IVF, primarily due to an increase in polyspermy (Fig. 2 D and F). Furthermore, the oocytes show significant lower developmental competence as the cleavage and blastocyst rate were all downregulated (Fig. 3). Collectively, these bidirectional perturbations-ranging from depletion to overexpression-establish JUNO as a dosage-dependent regulator of porcine sperm-egg recognition and fusion.

### Single cell proteomic reveals dynamic of proteins in porcine oocyte during fertilization

To identify key proteins associated with porcine sperm-oocyte fertilization, we analyzed differentially expressed proteins (DEPs) across pre- and post-fertilization. For each time point, 4 samples were examined, which were separated with each other (Fig. 4 A). A total of 499 proteins were found to be significantly altered in pairwise comparisons among the three stages (Fig. 4 B and Table S2). Compared to the MⅡ stage, 45 proteins were upregulated and 116 were downregulated at IVF-4h (Fig. 4 C). From IVF-4h to IVF-24h, 126 proteins showed increased abundance while 92 decreased (Fig. 4 D). All the upregulated and downregulated proteins across the three stages is summarized in Figure. 4 E and Table S3. These shifts in stage-specific and shared DEPs highlight pronounced remodeling of the proteome before and after fertilization.

**Figure 4.**
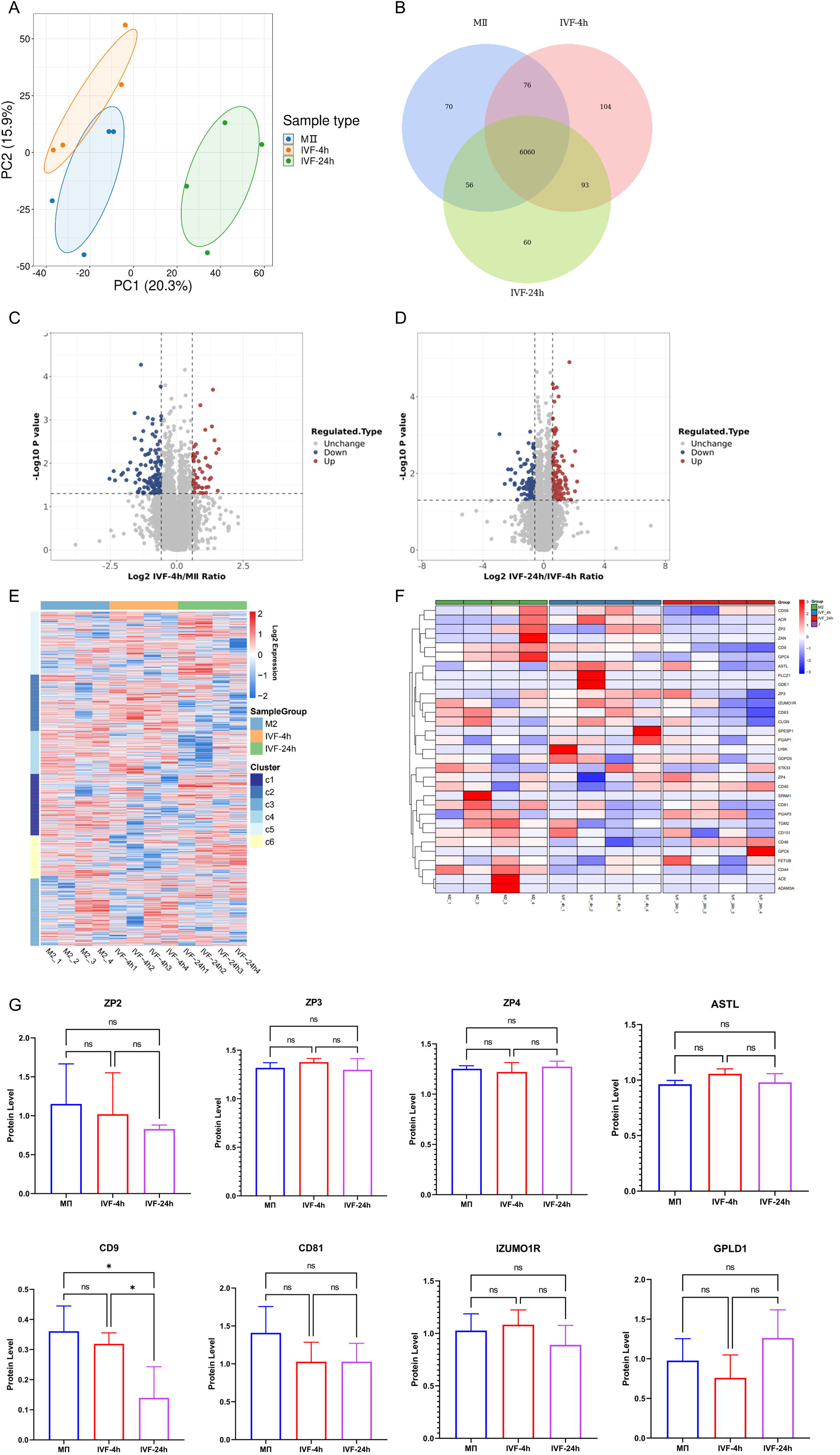
Proteomic profiling of porcine oocytes during the transition from fertilization to early embryogenesis. **(A)** Principal Component Analysis (PCA) of oocyte and embryo proteomes at MⅡ, IVF-4h, and IVF-24h stages. **(B)** Venn diagram showing the overlap of identified proteins across the three developmental stages. **(C, D)** Volcano plots of differentially expressed proteins (DEPs) comparing MⅡ vs. IVF-4h (C) and IVF-4h vs. IVF-24h (D). Red and blue points denote significantly up- and down-regulated proteins, respectively. **(E)** Hierarchical clustering heatmap of differentially expressed genes (DEGs) across fertilization stages. **(F)** Hierarchical clustering heatmap of DEPs; columns represent individual biological replicates and rows represent unique proteins. Color scale indicates normalized relative expression (red, high; blue, low). **(G)** Quantitative analysis of select fertilization-related proteins. Bar graphs show normalized abundance for zona pellucida proteins (ZP2, ZP3, and ZP4), metalloproteinase ASTL, tetraspanins CD81 and CD9, sperm receptor JUNO, and phospholipase GPLD1 across the indicated stages. Data are presented as mean ± SEM. Statistical significance was determined by one-way ANOVA. *P < 0.05; ns, not significant.

Given the enrichment of fertilization-related pathways, we next investigated proteins that have been reported to function in mammalian sperm–egg recognition. The proteomic analysis confirmed that porcine oocytes express ZP2, ZP3, and ZP4, while lacking ZP1 (Fig. 4 F and G), consistent with previous studies (Izquierdo-Rico et al., 2009). Following penetration of the zona pellucida, spermatozoa must recognize and fuse with oocyte membrane proteins. Among these, IZUMO1 on sperm and its oocyte receptor JUNO represent ligand-receptor pair demonstrated to be indispensable for gamete fusion. In porcine oocytes, JUNO was consistently detected at the MⅡ, IVF-4h, and IVF-24h stages (Fig. 4 F and G). Additional tetraspanins, including CD9 and CD81, were also present, aligning with their reported roles in human oocytes. Furthermore, early embryos, suggesting a broader contribution of GPI-anchored proteins to fertilization (Fig. 4 G and S1).

### Comparative profiling of membrane-associated proteins identifies potential drivers of the polyspermy block

To identify the molecular effectors governing gamete fusion and the subsequent block to polyspermy, we systematically interrogated the expression and evolutionary context of key oocyte proteins. While JUNO and its membrane-organizing partner CD81 remained stable in total abundance across the fertilization transition (Fig. 4 G), their functional impact is known to be dictated by rapid post-fusion redistribution. In murine models, JUNO is cleared from the surface within 40 minutes to establish a definitive membrane block (Bianchi et al., 2014); however, our proteomic profiling suggests that porcine oocytes maintain a robust reservoir of JUNO and associated tetraspanins, such as CD9, which exhibited an obvious decline during the same interval (Fig. 4 G). This sustained presence of fusion-essential receptors may explain the high intrinsic susceptibility of porcine oocytes to polyspermy.

A key feature of JUNO is its rapid release into the perivitelline space after fertilization, which prevents additional sperm from binding. Our proteomic data revealed that GPI-specific phospholipase D (GPLD1), which is an enzyme that cleaves the GPI anchor and releases GPI-anchored proteins from the plasma membrane and is expressed in oocytes at MⅡ, IVF-4h, and IVF-24h stages (Fig. 4 G). This finding raises the hypothesis that GPLD1 may facilitate polyspermy block by cleaving JUNO or related GPI-anchored proteins from oocyte. Subsequent functional perturbations are required to determine whether GPLD1 directly regulates JUNO localization during fertilization.

### GPLD1 exhibits dynamic expression and redistribution in oocyte during fertilization

To determine whether GPLD1 plays a critical role in porcine sperm-egg recognition and in preventing polyspermy, we conducted a comparative analysis of its expression patterns across oocytes and early embryos from different mammalian species. Firstly, publicly available transcriptomic data for human, mice and porcine (Yan et al., 2021; An et al., 2024; Zhi et al., 2022) were acquired, and the transcription pattern of GPLD1 were analyzed. As illustrated in Fig. 5 A-C, GPLD1 transcripts are detectable in oocytes and preimplantation embryos across multiple mammalian species, including human, mouse, and porcine. In human oocytes, *GPLD1* expression is relatively low prior to fertilization but increases markedly at the zygote stage, reaching peak levels at the 2-cell stage and then subsequently declining. By contrast, in mice, *GPLD1* expression follows a continuous upward trajectory after fertilization. In porcine oocytes and embryos, GPLD1 is present throughout the preimplantation period, although its abundance gradually decreases as development progresses.

**Figure 5.**
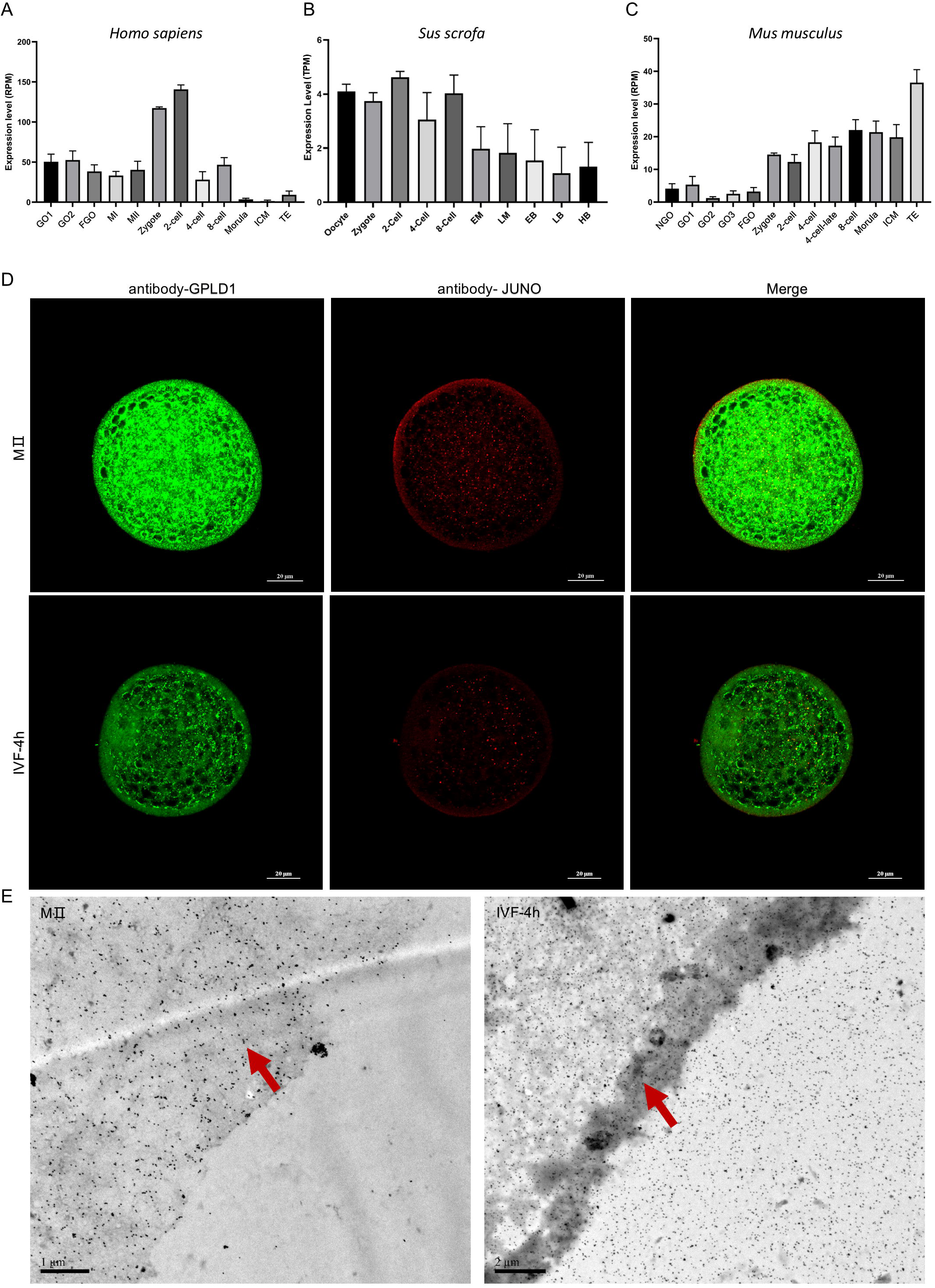
Expression and Subcellular Localization of GPLD1. **(A-C)** Expression profiles of GPLD1 across progressive developmental stages in *Homo sapiens* (A), *Sus scrofa* (B), and *Mus musculus* (C) models.**(D)** Co-immunofluorescence staining showing the spatial relationship between GPLD1 (green) and JUNO (red) on the oocyte membrane at MⅡ and IVF-4h. Scale bars, 20 μm. **(E)** Representative transmission electron microscopy (TEM) images showing the ultrastructural localization of GPLD1 on the membrane surface particles in porcine oocytes at the MⅡ and IVF-4h stages (red arrows indicate gold particles). Scale bars: 1 μm (MⅡ) and 2 μm (IVF-4h).

Next, the localization and redistribution of GPLD1 in porcine oocytes before and after fertilization were investigated. Proteomic profiling confirmed the presence of GPLD1 in MⅡ oocytes as well as in fertilized eggs at IVF-4h and IVF-24h post-insemination (Fig. 4 G). Immunofluorescence and transmission electron microscopy (TEM) analyses revealed that GPLD1 was diffusely distributed throughout the cytoplasm of unfertilized oocytes, but following fertilization it became enriched near the plasma membrane and was partially secreted into the perivitelline space (Fig. 5 D and E).

Consistent with these observed differences in spatiotemporal expression and localization patterns across species, we sought to determine if such functional divergence was mirrored at the primary sequence level. Beyond JUNO, we examined the sequence characteristics of GPLD1 across five mammalian species to assess its evolutionary variability (Fig. S1 D). Our comparative analysis reveals that GPLD1 exhibits significant interspecies sequence divergence, matching the heterologous expression profiles previously noted. Specifically, the sequence identity between pig and mouse GPLD1 is 75.03% (Fig. S1 E). Multiple sequence alignment further highlights numerous non-conserved regions and specific amino acid substitutions distributed throughout the protein (Fig. S1 D). This molecular diversity, coupled with the distinct expression trajectories observed in Fig. 5A-C, suggests that GPLD1 may have undergone species-specific functional adaptations to meet the unique requirements of the fertilization process in different organisms.

### GPLD1 enzymatic activity is essential for limiting sperm binding and ensuring monospermic fertilization

To determine the functional significance of GPLD1 during fertilization, we first employed an siRNA-mediated knockdown strategy in porcine oocytes. GPLD1-specific siRNA was microinjected into the oocyte cytoplasm at 32 h of *in vitro* maturation, whereas a non-targeting scrambled siRNA was used as a control. Subsequent immunofluorescence analysis confirmed the efficacy of the knockdown, with a significant reduction in GPLD1 fluorescence intensity observed across all developmental stages compared to the control group (Fig. 8 A-C). Membrane-localized JUNO was slowly cleared following sperm–egg fusion, becoming significantly attenuated by IVF-12h in oocytes from control group. In contrast, oocytes deficient in GPLD1 exhibited a striking retention of JUNO on the oocyte at both 4 h and 12 h post-insemination (Fig. 8 A-C). These results indicate that the timely depletion of JUNO from the zygote surface is strictly dependent on the presence of functional GPLD1.

Next, the outcome of fertilization competence was examined. Quantitative analyses demonstrated that loss of GPLD1 resulted in a significant increase in sperm binding to the oocyte surface at 4 h post-fertilization (Fig. 6 A and C). By contrast, assessment at 12 h post-fertilization revealed that GPLD1-deficient oocytes exhibited a markedly reduced capacity for normal double-pronuclear (2PN) formation (Fig. 6 B and D). Furthermore, by effectively mitigating polyspermy, these oocytes demonstrated significantly higher developmental competence, as both cleavage and blastocyst formation rates were upregulated (Fig. 7). Together, these observations indicate that GPLD1 plays a dynamic role during fertilization and that its depletion disrupts the coordination between initial sperm–oocyte interactions and subsequent pronuclear development.

**Figure 6.**
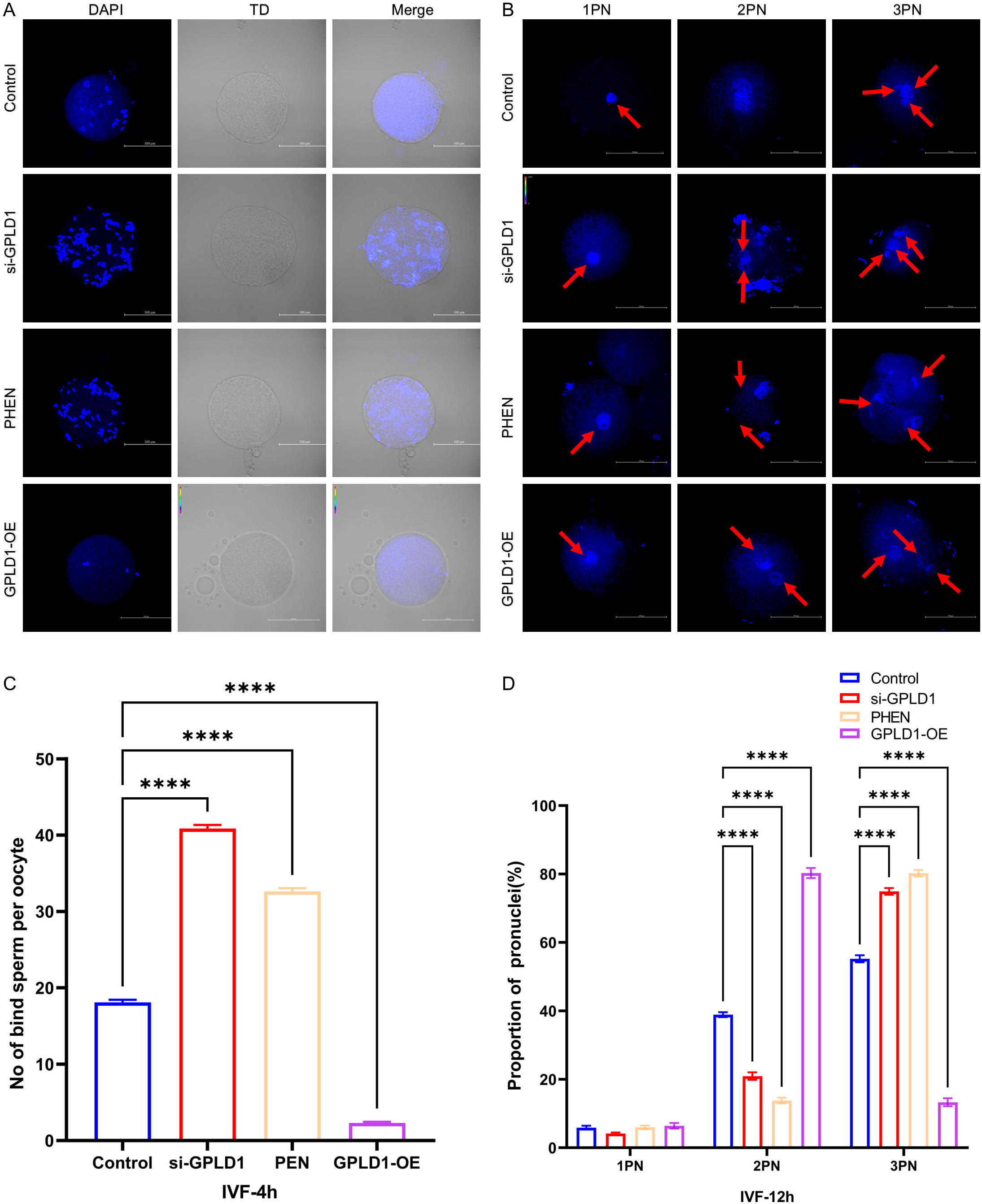
GPLD1 Regulates Sperm Binding and Blocks Polyspermy. **(A)** Representative fluorescence images showing sperm binding to porcine oocytes under different treatments (Control, si-GPLD1, PHEN, and GPLD1-OE) at IVF-4h. Oocytes were stained with DAPI (blue) to visualize sperm nuclei. Scale bars, 100 μm. **(B)** Representative images of pronuclear formation in fertilized oocytes at 12 h post-IVF under different treatments. Arrows indicate pronuclei. Oocytes exhibiting one pronucleus (1PN), two pronuclei (2PN), or three pronuclei (3PN) were observed, reflecting variations in fertilization and polyspermy among the groups. Scale bars, 100 μm. **(C)** Bar chart comparing the number of bound sperm per oocyte at IVF-4h across Control, si-GPLD1, PHEN (GPLD1 inhibitor), and GPLD1-OE (overexpression) groups. Data are presented as mean ± SEM; ****P < 0.0001.**(D)** Proportion of pronuclei (1PN, 2PN, and polyspermic 3PN) observed at IVF-12h across the different experimental groups. Data are presented as mean ± SEM; ****P < 0.0001.

**Figure 7.**
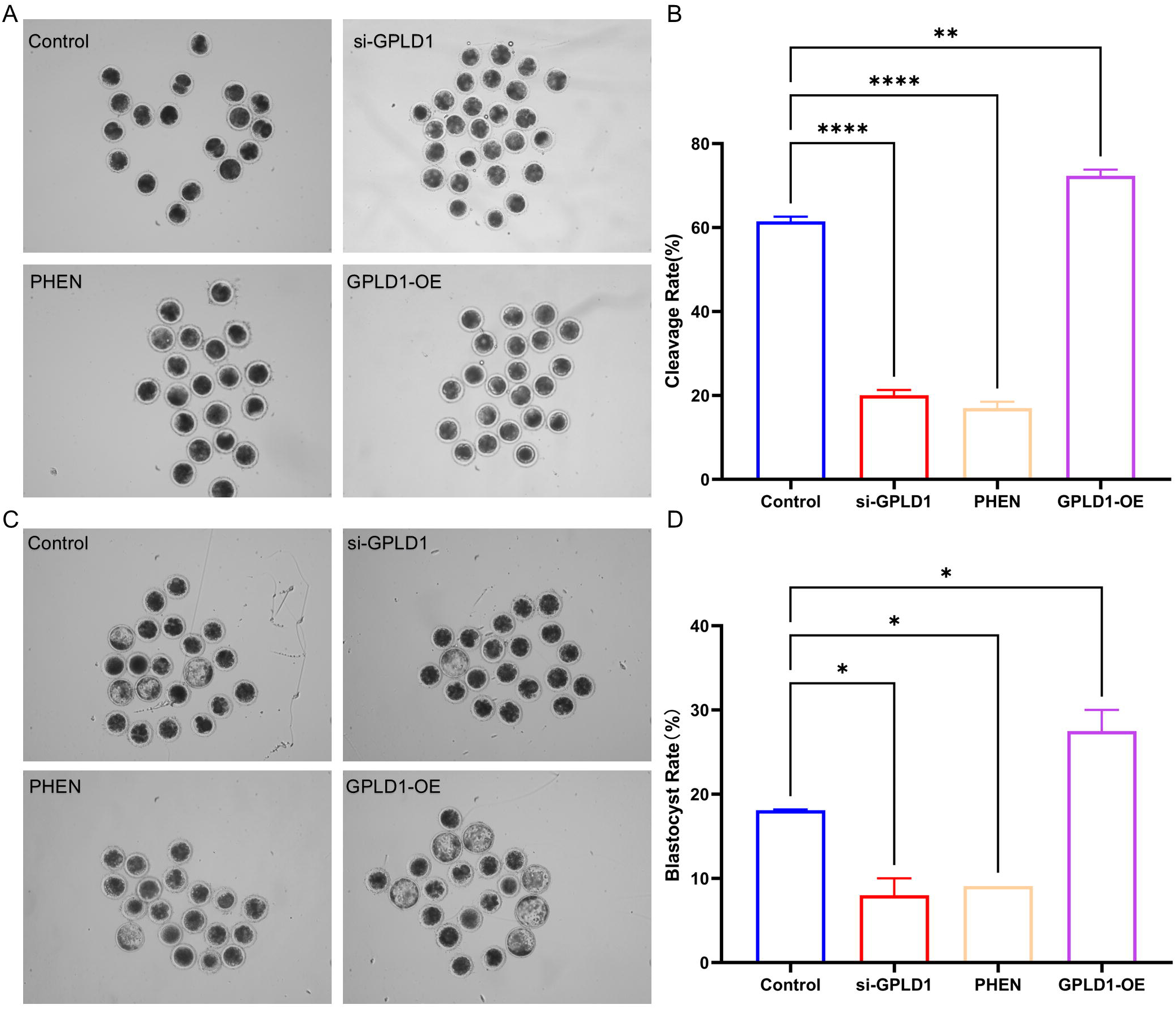
Effect of GPLD1 expression levels on early embryonic development. **(A)** Representative images of embryos at the cleavage stage. Micrographs showing the morphology of embryos in different treatment groups (Control, si-GPLD1, PHEN, and GPLD1-OE) during early cleavage. **(B)** Comparison of cleavage rates across groups. Statistical analysis of the percentage of embryos reaching the cleavage stage. Data are presented as mean ± SEM. **P < 0.01, ****P < 0.0001. **(C)** Representative images of embryos at the blastocyst stage. Micrographs illustrating the development of embryos into blastocysts in the four experimental groups. **(D)** Comparison of blastocyst rates across groups. Statistical analysis of the percentage of embryos reaching the blastocyst stage. Data are presented as mean ± SEM. *P < 0.05.

### GPLD1 enzymatic activity is required for JUNO shedding during fertilization

To further delineate the mechanistic basis of GPLD1 function, with particular emphasis on its enzymatic activity in regulating JUNO dynamics, we next pharmacologically inhibited GPLD1 using 1,10-phenanthroline (PHEN), a well-characterized metalloprotease inhibitor known to suppress GPLD1 catalytic activity (Mann and Sevlever, 2001). Notably, and in agreement with the knockdown phenotype, PHEN-treated oocytes exhibited a significant increase in sperm binding to the oocyte at 4 h post-fertilization (Fig. 6 A, C and S2 A, B). Moreover, these oocytes displayed a markedly reduced capacity for 2PN formation at 12 h post-fertilization (Fig. 6 A and C). Furthermore, by effectively mitigating polyspermy, these oocytes demonstrated significantly higher developmental competence, as both cleavage and blastocyst formation rates were upregulated (Fig. 7). Immunofluorescence staining and confocal microscopy revealed that the surface-localized JUNO signal (red) was rapidly depleted following insemination, becoming nearly undetectable by 12 h post-fertilization in control group (Fig. 8 A). This loss of JUNO was accompanied by a concomitant reduction in GPLD1 fluorescence intensity (green) as fertilization progressed (Fig.8, A and C). In contrast, JUNO remained in the PHEN-treated oocyte at both 4 h and 12 h post-IVF, exhibiting fluorescence intensities significantly higher than those of the control group (Fig. 8 A and B). Quantitative analysis confirmed that the persistence of JUNO directly correlated with the successful inhibition of GPLD1, as evidenced by the dramatic reduction in GPLD1 protein levels across all stages in the PHEN group (Fig. 8 C).

**Figure 8.**
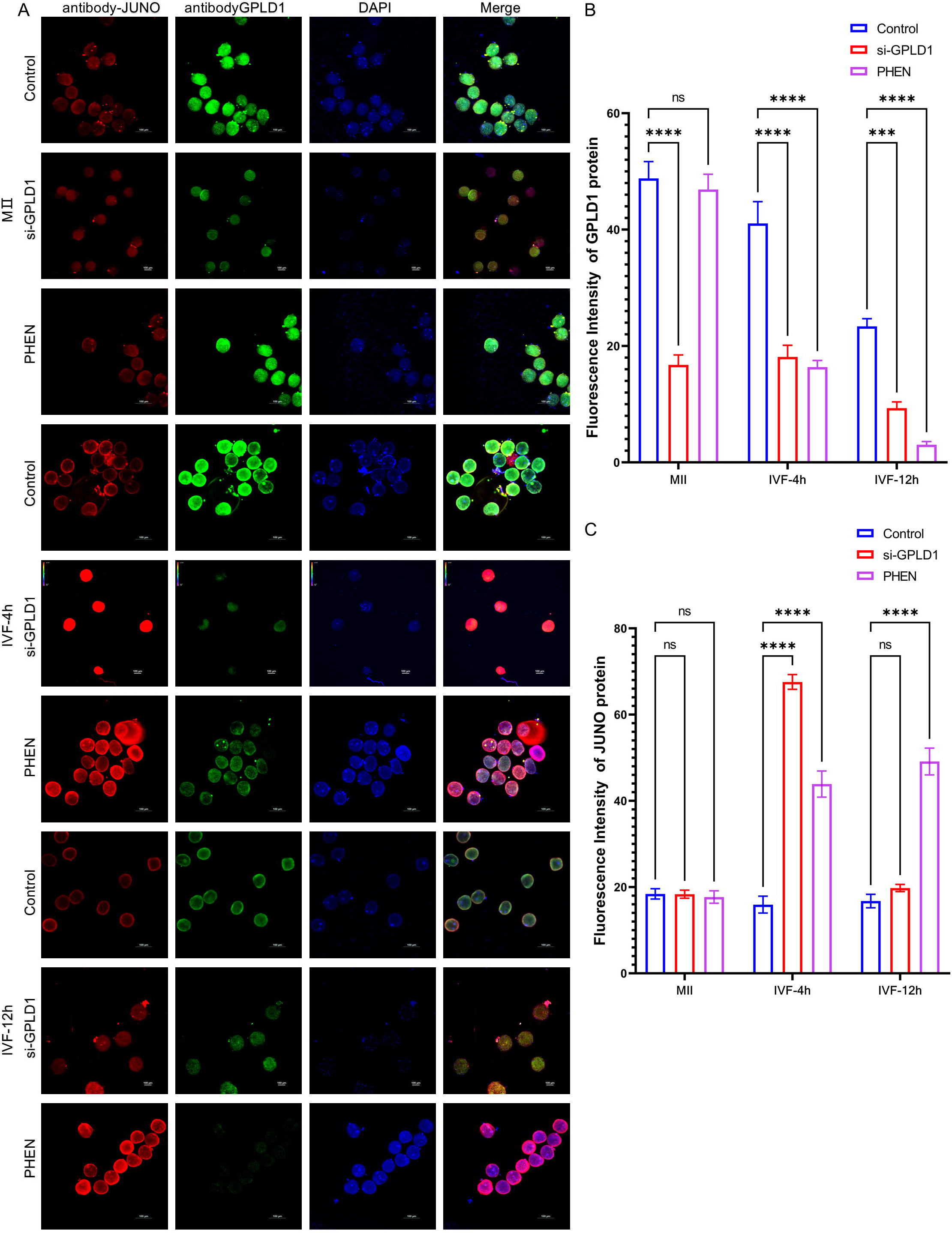
GPLD1 regulates the fluorescence intensity and localization of JUNO protein during in vitro fertilization. **(A)** Immunofluorescence staining of JUNO and GPLD1. Representative confocal images showing JUNO (red), GPLD1 (green), and DAPI (blue, nuclei) in control group, si-GPLD1 and PHEN groups across three stages: MⅡ oocytes, IVF-4h, and IVF-12h. Scale bars are indicated in the images. Scale bars, 100 μm. **(B)** Quantitative analysis of JUNO protein fluorescence intensity. Statistical comparison shows that si-GPLD1 and PHEN significantly increases the retention of JUNO protein on the oocyte/embryo surface at IVF-4h and IVF-12h compared to Control group. Data are presented as mean ± SEM. ****. P < 0.0001. ns, not significant. **(C)** Quantitative analysis of GPLD1 protein fluorescence intensity. The significant reduction in GPLD1 fluorescence in the si-GPLD1 and PHEN group across all stages validates the efficiency of the siRNA-mediated knockdown. Data are presented as mean ± SEM. ****P < 0.0001. ns, not significant.

Finally, to determine whether GPLD1 abundance represents an essential factor for fertilization, we examined whether the elevation of GPLD1 expression in porcine oocytes is sufficient to diminish sperm recruitment. In contrast to the siRNA and pharmacological inhibition approaches, oocytes overexpressing GPLD1 exhibited a dramatic reduction in sperm binding at 4 h post-insemination compared with that in control group (Fig. 6 A and C). At 12 h post-fertilization, an increase in the proportion of monospermic (2PN) zygotes, accompanied by a decrease in multi-pronuclear formation (Fig. 6 B and D). Furthermore, by overexpressing GPLD1, these oocytes demonstrated significantly higher developmental competence, as both cleavage and blastocyst formation rates were elevated (Fig. 7). These findings indicate that GPLD1 negatively regulate sperm adherence while promoting the monospermic fertilization.

### GPLD1 mediates enzymatic cleavage of JUNO in porcine oocyte

To track membrane protein dynamics and investigate whether this spatiotemporal coordination reflects a functional necessity, the enzymatic requirement of GPLD1 in JUNO shedding was verified through gain-of-function and pharmacological inhibition assays. Overexpression of GPLD1 was sufficient to trigger a dramatic reduction in surface JUNO abundance compared to controls (Fig. 9 A, B and S3 A, B). Conversely, treatment with PHEN effectively abrogated the fertilization-induced membrane recruitment of GPLD1 and impaired JUNO–enhanced Cyan Fluorescent Protein (JUNO–eCFP) depletion, leading to its persistent retention on the oocyte (Fig. 9 A and B). These results establish GPLD1 as the requisite enzymatic driver of JUNO cleavage.

**Figure 9.**
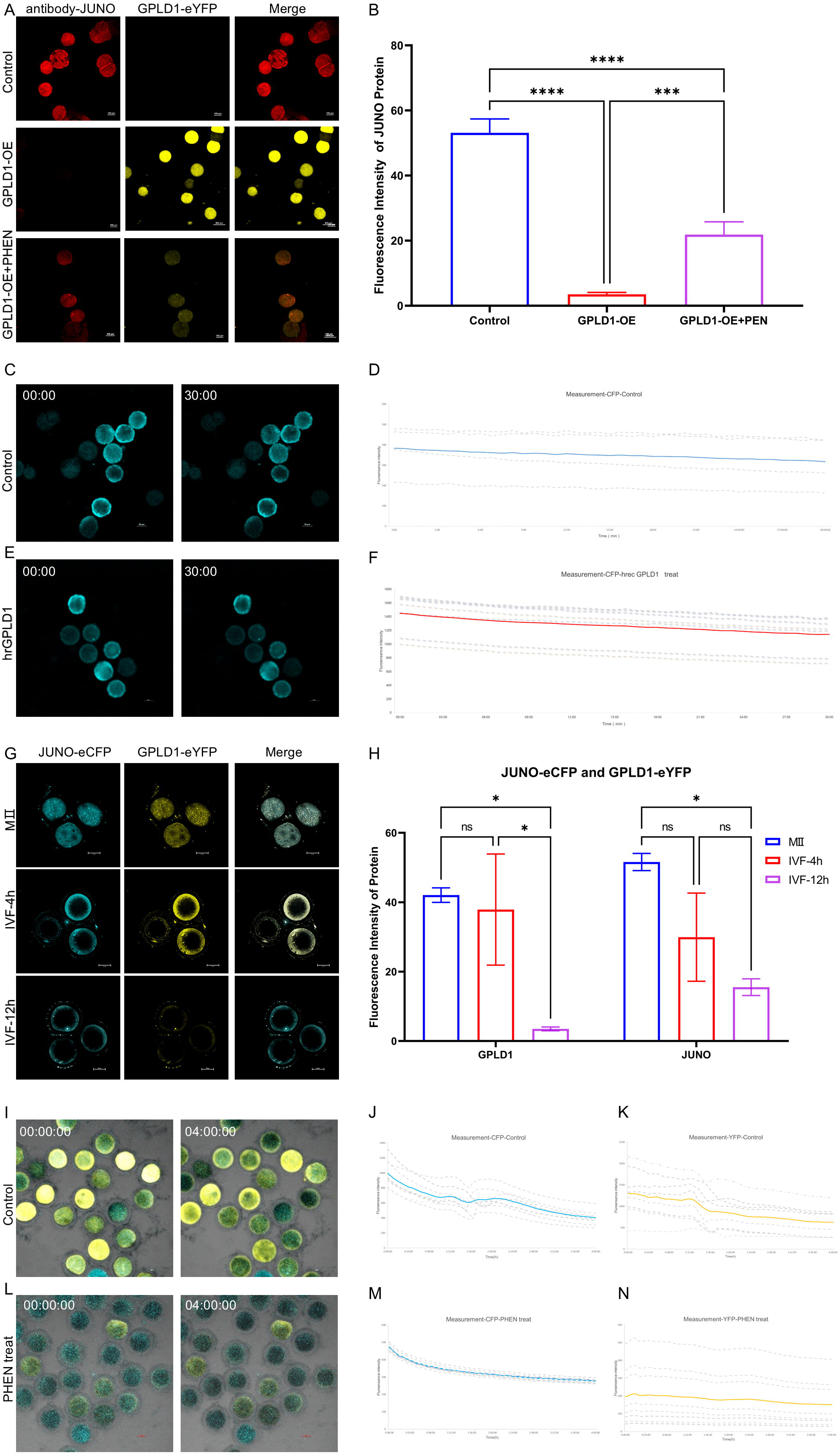
GPLD1 directs the enzymatic shedding of JUNO from the porcine oocyte during fertilization. **(A)** Immunofluorescence analysis of JUNO (red) in oocytes overexpressing GPLD1-eYFP (yellow) with or without PHEN and Control. Scale bars, 100 μm. **(B)** Quantification of JUNO fluorescence intensity from (A). Data represent mean ± SEM. Statistical significance was determined by one-way ANOVA. Data are presented as mean ± SEM. ∗∗∗P<0.001, ∗∗∗∗P<0.0001. **(C, E)**Live-cell time-lapse imaging of JUNO-eCFP (Control) and JUNO-ECFP+ human recombinant GPLD1 protein (hrGPLD1) shedding over 30 min. Scale bars, 50 μm. **(D, F)** Depletion of JUNO-eCFP following the application of exogenous hrGPLD1 and Control. Bottom panels show individual intensity traces over time for multiple oocytes. Time is indicated in minutes. **(G)** Representative images of oocytes co-injected with JUNO-eCFP and GPLD1-eYFP showing their dynamic localization during IVF. **(H)** Quantification of JUNO-eCFP and GPLD1-eYFP fluorescence intensities at MII, IVF-4h, and IVF-12h stages. Data are presented as mean ± SEM. *P < 0.05. ns, not significant. **(I)** Representative time-lapse imaging of oocytes co-expressing JUNO-eCFP (cyan) and GPLD1-eYFP (yellow) during in vitro fertilization. (See also Video 3). **(J, K)** Quantitative analysis of JUNO-eCFP (J) and GPLD1-eYFP (K) fluorescence intensity over time in control zygotes group. **(L)** Time-lapse imaging of IVF oocytes treated with PHEN. (See also Video 4). **(M)** Quantitative analysis of JUNO-eCFP (M) and GPLD1-eYFP **(N)** fluorescence intensity in PHEN-treated zygotes.

To investigate the direct effect of GPLD1 on JUNO protein dynamics, we first performed live-cell confocal imaging on oocytes expressing JUNO-eCFP. Upon the addition of human recombinant GPLD1 protein (hrGPLD1), a rapid depletion of JUNO-eCFP from the oocyte was observed within 30 min (Fig. 9 C and E). Quantitative analysis of the fluorescence intensity of JUNO-eCFP ratio showed a significant decrease, confirming that GPLD1-mediated cleavage is the primary driver for JUNO release into the perivitelline space (Fig. 9 D and F; and Video 1, 2).

### Live-cell imaging reveals that JUNO shedding is driven by the transient spatiotemporal recruitment of GPLD1

To elucidate the precise kinetic characteristics of JUNO shedding during fertilization and its spatiotemporal coordination with GPLD1, we performed live-cell time-lapse imaging of porcine oocytes co-expressing JUNO-eCFP and GPLD1-enhanced Yellow Fluorescent Protein (GPLD1-eYFP).Live-cell imaging showed synchronized dynamics of JUNO and GPLD1 during fertilization in oocytes from control group. JUNO transiently increased after sperm fusion a from the oocyte (Fig. 9 G, H and S4 A, B). In parallel, GPLD1 was briefly recruited to the membrane before dispersing into the cytoplasm, tightly coupling with JUNO shedding. In fertilized oocytes from control group, these two proteins exhibited highly synchronized and distinct dynamic localization patterns related to sperm-oocyte fusion (Video 3). Following the onset of fusion, the oocyte JUNO-eCFP signal did not immediately diminish; instead, it displayed a significant, transient upward spike (Fig. 9 I-K). Subsequently, as membrane-recruited GPLD1 catalyzed the enzymatic cleavage of its substrate, the JUNO-eCFP signal underwent a progressive and irreversible decay (Fig. 9 I-K; and Video 3). Coincident with the depletion of surface JUNO, the oocyte GPLD1-eYFP signal subsided, with the protein dissociating from the membrane to revert to a diffuse cytoplasmic distribution (Fig. 9 I-K).

To validate the functional necessity of this spatiotemporal recruitment, we employed pharmacological intervention with PHEN. Live-cell imaging and quantitative fluorescence analysis demonstrated that PHEN treatment completely abolished this spatiotemporal coordination (Video 4). Specifically, PHEN significantly inhibited the pulsed recruitment of GPLD1-eYFP, restricting its membrane fluorescence to baseline levels (Figure. 9, L-N). As a direct consequence, the membrane clearance of JUNO-eCFP was completely halted. Rather than exhibiting the biphasic kinetics observed in oocytes from control group, JUNO remained continuously and stably retained in porcine oocytes post-fertilization. Together, these kinetic and pharmacological data conclusively demonstrate that fertilization-induced JUNO cleavage is not a passive internalization event, but a precisely regulated spatial process critically dependent on the transient membrane recruitment and enzymatic activation of GPLD1.

## DISCUSSION

Sperm-egg recognition and fusion is essential for the success of animal fertilization, and the high rate of polyspermy in pig eggs results in more abnormal early embryonic development and lower litter size. JUNO mediate animal sperm-egg specific recognition, and is detached from the oocyte membrane after fertilization, preventing extra sperm from entering the egg. In the present study, we integrated proteomics with targeted functional analyses to characterize the molecular determinants of gamete recognition in the porcine oocyte. Our findings delineate a regulatory mechanism wherein the dynamic remodeling of membrane-associated proteins-specifically the GPI-anchored receptor JUNO and its enzymatic regulator GPLD1, sperm–egg interactions and establishes a block to polyspermy. By determining the enzymatic activity of GPLD1 during fertilization, we demonstrate its critical contribution to restricting sperm adherence and safeguarding monospermic fertilization.

### The expression and function of JUNO in regulating porcine oocyte fertilization

JUNO serves as the essential oocyte cytoplasm membrane receptor that binds the sperm protein IZUMO1 to ensure the success of fertilization. In mice, JUNO becomes undetectable on the oocyte surface within approximately 40 minutes post-fertilization, a timeframe that precisely aligns with the establishment of the membrane block to prevent extra sperm recognition (Bianchi et al., 2014). The analyses in current study reveal that the JUNO in porcine oocytes declined post fertilization, but remain at a detectable level through early cleavage stages. The observation with TEM also showed that JUNO is partially released to PVS post fertilization, and the knockdown of JUNO or PIPLC treatment significantly disrupted porcine sperm-oocyte fusion. Porcine oocyte is exceptional susceptibility to polyspermy during *in vitro* fertilization compared to other mammalian models (Himaki et al., 2025). The sustained presence of JUNO in porcine oocyte post-fertilization may allow the embryo to retain sperm-binding capacity, thereby contributing to the high rate of polyspermy observed in porcine oocyte during fertilization.

### Regulation of porcine sperm-oocyte recognition by the enzymatic cleavage GPI-anchored proteins

JUNO belongs to the folate receptor family but is unique in its function as a glycosylphosphatidylinositol-anchored protein (GPI-AP) for molecule adhesion rather than a transporter (Chen et al., 2013; Wibowo et al., 2013). The essential role of GPI-APs in mammalian fertilization has been recognized since the landmark discovery of JUNO (Bianchi E et al., 2014), yet the molecule governing their rapid post-fertilization cleavage remains a fundamental question in reproductive biology. A distinctive characteristic of GPI-APs is their susceptibility to cleavage by specific phospholipases, which facilitates the release of the protein from the cell membrane. By proteomic analyses and IF observation, we identified the existence of GPLD1 (glycosylphosphatidylinositol specific phospholipase D1) in porcine oocyte. GPLD1 is a soluble enzyme that plays a key role in regulating cell-surface signaling and function by specifically cleaving GPI anchors (Cao et al., 2023). Unlike other phospholipases such as PIPLC and GPI-PLC, GPLD1 uniquely hydrolyzes GPI anchors that contain acylated inositol residues (Raikwar et al., 2005). In mice, its catalytic activity depends critically on conserved histidine residues at positions 29, 125, 133, and 158 within the active site (Raikwar et al., 2005). By cleaving GPI anchors, GPLD1 releases membrane-bound GPI-APs from the cell surface, thereby modulating their localization and function (Fujihara and Ikawa, 2016). To test the function of GPLD1, its expression in porcine oocytes was knocked down by microinjection of specific siRNA or treatment with an inhibitor (1-10-phenanthroline). This treatment led to an elevated level of JUNO in porcine oocytes and improved their *in vitro* fertilization outcomes. Meanwhile, the overexpression of GPLD1 reduced the level of JUNO and impaired fertilization in porcine oocytes. Notably, GPLD1 is significantly upregulated in oocytes derived from large follicles and persists during early embryonic development (Gad et al., 2020; Zhi et al., 2022), suggesting a potential role in oocyte maturation, fertilization and embryo development. Based on these findings, it can be concluded that GPLD1 regulates JUNO activity and fertilization efficiency in porcine oocytes.

### GPLD1 Regulates JUNO Dynamic in Porcine Oocyte During Fertilization

Given its ability to cleave GPI-anchored proteins, GPLD1 is positioned to contribute to promote the shedding or redistribution of key membrane components. Such regulation may reduce the binding capacity of additional sperm and facilitate the establishment of the membrane block to polyspermy. Thus, we propose that GPLD1 enzymatically cleaving the GPI anchor of JUNO to drive its rapid shedding into the perivitelline space. By eliminating the physical receptor required for IZUMO1-mediated binding, the oocyte effectively disarms its surface receptivity, thereby terminating further sperm-egg interactions at the level of the plasma membrane.

Further research with live-cell imaging, we observed that hrGPLD1 treatment induce a sharp decline of JUNO-eCFP, which provide compelling evidence for the GPLD1 and mediated cleavage of JUNO. Then, GPLD1-eYFP and JUNO-eCFP redistributed to the aera close to the membrane and their protein level both reduced at 12h post-fertilization. When those porcine oocytes were incubated sperm, the timelapse observation reveal that PHEN treatment could slow down the fluorescence intensity decline especially for JUNO-eCFP. Based on above evidence, GPLD1 serves as the GPI anchor hydrolase in porcine oocyte to cleavage JUNO following primary sperm-oocyte recognition and fusion to prevent extra sperm entry.

### Species-specific sperm-oocyte recognition zona pellucida proteins in porcine oocytes

Besides sperm-oocyte recognition mediated by membrane protein, sperm firstly recognizes and penetrates the zona pellucida-a glycoprotein-rich extracellular matrix. Our proteomic analysis revealed that the porcine ZP is composed of ZP2, ZP3, and ZP4, notably lacking ZP1. Unlike mice, where ZP1 provides critical structural cross-linking for ZP filaments, porcine oocytes appear to rely exclusively on ZP2, ZP3, and ZP4 to establish a functional sperm-binding scaffold (Nishio et al., 2024; Yonezawa et al., 2012). Among these, ZP3 and ZP4 are recognized as the principal molecular interfaces mediating primary sperm recognition and adherence (Yonezawa et al., 2012). The ZP block acts as a macroscopic mechanical filter which relies on the exocytosis of cortical granules (CGs) to trigger the enzymatic cleavage of ZP2 by ovastacin (ASTL), thereby hardening the matrix and preventing further sperm penetration (Burkart et al., 2012). In current study, the level of ASTL did not change significantly, thus further research is required to elucidate its function in cleavage ZP proteins in porcine oocyte post fertilization. These findings reinforce the notion that ZP composition has evolved divergently across mammals, likely reflecting distinct sperm-oocyte recognition and susceptibilities to polyspermy.

### Concluding remarks

Fertilization is a highly coordinated, multistep process that relies on precise molecular interactions between sperm and oocyte, beginning with ZP recognition and culminating in sperm-egg membrane fusion and activation of polyspermy block mechanisms. While the requirement for the GPI-anchored receptor JUNO in primary sperm-egg recognition is well documented, the spatiotemporal mechanisms governing its rapid post-fertilization cleavage have remained an enduring question in reproductive cell biology. In this study, we provide compelling evidence that GPLD1 is the essential enzymatic factor responsible for JUNO shedding in porcine oocyte. Our findings establish a continuous regulatory axis in which GPLD1-mediated cleavage of JUNO following gamete species specific recognition and safeguards subsequent embryogenesis. The physiological indispensability of the GPLD1-mediated membrane JUNO shedding, highlighting its potential as a biomarker or therapeutic target for optimizing assisted reproductive technologies (ART) in for animals.

## Materials and methods

### Chemicals

All chemicals used in this study were purchased from Merck KGaA (Darmstadt, Germany and/or its affiliates) unless otherwise indicated.

### Collection of Porcine Oocytes, In Vitro Fertilization, and Embryo Culture

Porcine cumulus–oocyte complexes (COCs) were recovered and rinsed in HEPES-buffered Tyrode’s lactate (TL-HEPES) medium. COCs were subsequently cultured in maturation medium (TCM-199 supplemented with cysteine, glucose, sodium pyruvate, EGF, LH, FSH, polyvinyl alcohol, and antibiotics) at 38.5 °C in a humidified atmosphere of 5% CO_2_. After 44 h of maturation, cumulus cells were removed by incubation with 0.1% (w/v) hyaluronidase. Only denuded oocytes exhibiting an intact plasma membrane, homogeneous cytoplasm, and a visible first polar body (PB1) were selected for further experiments.

Following maturation, oocytes were rinsed and equilibrated in modified Tris-buffered medium (mTBM). For insemination, liquid boar semen (Huaxia Su Bu Biotechnology Co., Ltd.) was processed via centrifugation (1900xg for 4 min) and resuspended in mTBM to the required concentration. Oocytes were then co-cultured with sperm at a ratio of 500:1 in mTBM droplets. Fertilization was maintained at 38.5 °C in a humidified incubator with 5% CO₂. Approximately 4–6 h post-insemination, zygotes were removed from the fertilization medium and transferred into 30 mm culture dishes containing HEPES-buffered Tyrode’s lactate (TL-HEPES) solution. Adherent spermatozoa were removed from the zona pellucida by using pipette. Zygotes were subsequently washed three times in Porcine Zygote Medium-3 (PZM-3) before being transferred into 50 µL developmental culture droplets. All embryos were cultured at 38.5 °C in a humidified atmosphere of 5% CO_2_ and 100% humidity.

### Single-Cell Proteomics

Single-Cell Proteomics was performed according to previous study (Jiana Huang et al., 2023). Briefly, porcine oocyte samples were rinsed three times on ice and lysed in 8 M urea buffer containing 1% protease inhibitor cocktail using a high-intensity ultrasonic processor (Scientz). Lysates were cleared by centrifugation (12,000 × g, 10 min, 4 °C), and protein concentration in the supernatant was quantified via BCA assay according to the manufacturer’s instructions. For digestion, proteins were reduced with 5 mM dithiothreitol (30 min, 56 °C) and alkylated with 11 mM iodoacetamide (15 min, room temperature, in the dark). Urea was diluted to < 2 M with 200 mM TEAB, and proteins were digested sequentially with trypsin at 1:50 (overnight) and 1:100 (4 h) enzyme-to-protein ratios. Resulting peptides were desalted using Strata™ X solid-phase extraction columns.

Peptides were reconstituted in solvent A (0.1% formic acid, 2% acetonitrile in water) and separated on a home-made C18 reversed-phase column (25 cm × 100 μm i.d.) using a NanoElute UHPLC system (Bruker Daltonics) at 500 nL/min. The gradient profile was: 6-24% solvent B (0.1% formic acid in acetonitrile) over 0-14 min; 24-35% B over 14-16 min; 35-80% B over 16-18 min; and 80% B held from 18-20 min. Eluted peptides were analyzed on a timsTOF Pro 2 mass spectrometer equipped with a nano-electrospray source (1.75 kV), operating in dia-PASEF mode. MS/MS spectra were acquired over m/z 400–850 with 20 PASEF frames per cycle; precursors with charge states 0–5 were selected for fragmentation.

Raw files were processed in DIA-NN (v1.8) against the *Sus scrofa* reference proteome (*Sus_scrofa*_9823_PR_20231009.fasta; 46,179 entries) concatenated with a reverse decoy database. Trypsin/P specificity was applied with up to 1.0 missed cleavages. Carbamidomethyl (Cys) was set as a fixed modification. Precursor and fragment mass tolerances were set to 5 ppm and 0.02 Da, respectively. All results were filtered to a 1% false discovery rate (FDR) at the peptide and protein levels. To understand the functional characteristics of different proteins, comprehensive functional annotation on the identified proteins (Table s1) was performed across multiple dimensions, including Gene Ontology (GO), Protein domains, KEGG pathways, COG/KOG functional classification, Subcellular localization, Reactome, WikiPathways, Hallmark, and Transcription factors (TF).

The amino acid sequences of JUNO and GPLD1 from five mammalian species including mouse (Mus musculus), human (Homo sapiens), pig (Sus scrofa), sheep (Ovis aries), and cattle (Bos taurus)—were retrieved from the UniProtKB database (https://www.uniprot.org/). Multiple sequence alignments (MSA) were performed using ClustalW (used MUSCLE) and further processed using uniprot to visualize conserved residues. Identical and partially conserved residues were highlighted based on their level of conservation. Potential N-glycosylation sites and GPI-anchor signal sequences were predicted using UniProt annotations.To quantify the evolutionary distance between species, pairwise sequence identity matrices were generated. The percentage of identity for each species pair was calculated through global alignment.

### PIPLC and PHEN treatments

To enzymatically remove GPI-anchored proteins, phosphatidylinositol-specific phospholipase C (PIPLC; Sigma-Aldrich) was prepared as a stock solution in TL-HEPES and diluted in culture medium to a final working concentration of 0.2 U/mL (ensuring the final TL-HEPES concentration remained below 0.1% v/v). Zona-free MⅡ oocytes, previously transfected with a JUNO expression construct, were incubated with 0.2 U/mL PIPLC for 1 h at 37.5 °C. Following treatment, oocytes were rinsed in TL-HEPES and subjected to either in vitro fertilization or immunofluorescence analysis.

To inhibit metalloproteinase activity, oocytes were treated with 1,10-Phenanthroline (PHEN; Sigma-Aldrich). PHEN was dissolved in TL-HEPES and diluted in culture medium to the specified working concentrations (final TL-HEPES < 0.1% v/v). Zona-free MⅡ oocytes were incubated with PHEN for 1 h at 37.5 °C, followed by extensive washing in TL-HEPES prior to subsequent experimental procedures.

### siRNA Microinjection and Knockdown

The mRNA sequences for porcine JUNO (GenBank accession no. XM_013979283) and GPLD1 (GenBank accession nos. XM_013977471 and XM_001925909) were retrieved from GenBank. Small interfering RNAs (siRNAs) targeting these genes, as well as a non-targeting negative control (scrambled) siRNA, were synthesized by GenePharma (Shanghai, China). The specific siRNA sequences are provided in Table 1. To achieve gene knockdown, MⅡ oocytes were denuded 32 h after the initiation of maturation by incubation in 0.1% (w/v) hyaluronidase and subsequent rinsing in TL-HEPES. siRNAs were prepared as 75 µM stock solutions in RNase-free water. Approximately 5–10 pL of the siRNA solution was microinjected into the cytoplasm of the oocytes using a microinjector. Following injection, oocytes were cultured in IVM medium for an additional 12 h to allow for protein depletion and completion of maturation. Following the 12h incubation, a subset of MⅡ oocytes underwent zona pellucida removal and were subjected to IVF. Resulting zygotes were fixed at 4 h and 12 h post-insemination for downstream immunofluorescence and expression analysis.

**Table 1.**
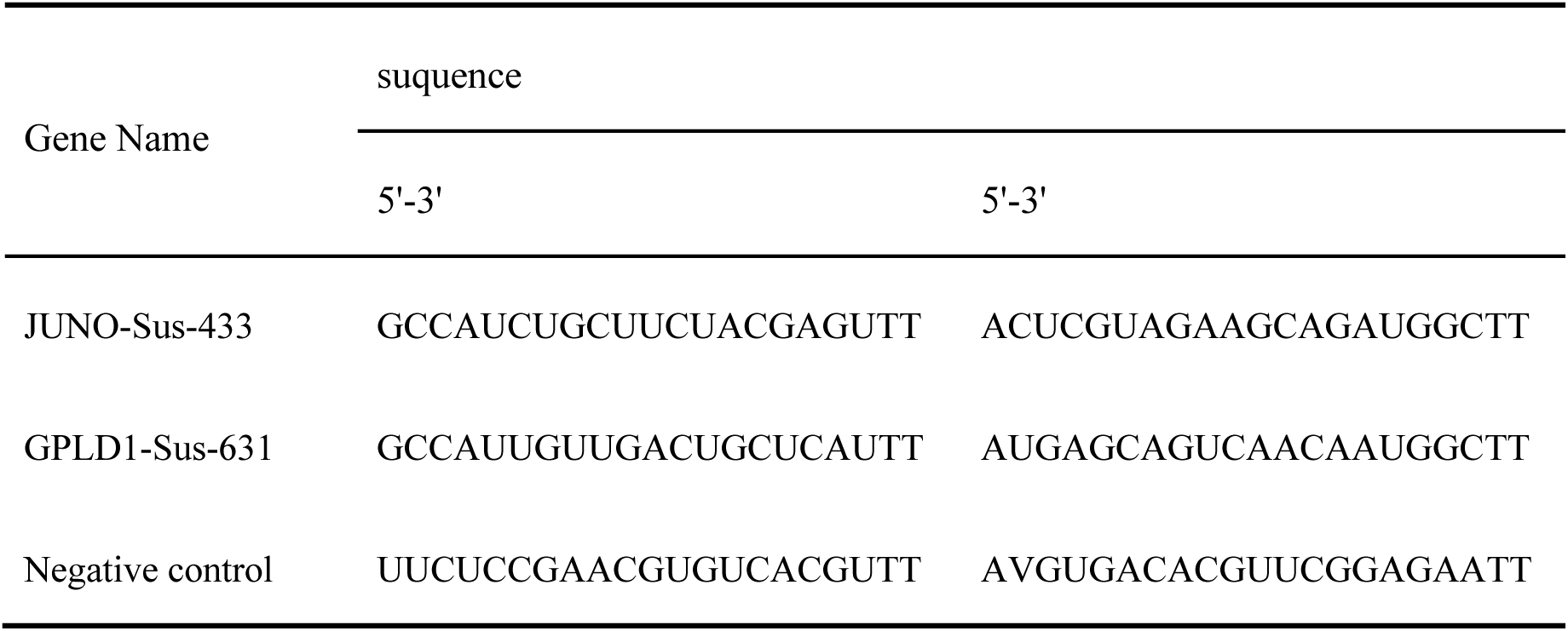
siRNA sequence.

### mRNA Microinjection and Functional Analysis

The capped and polyadenylated mRNA encoding porcine JUNO-eCFP or/and GPLD1-eYFP was synthesized by (GenScript Corporation, Nanjing, China). The mRNA sequence was optimized for mammalian expression and purified via HPLC to ensure high translational efficiency and minimal immunogenicity. Upon receipt, the lyophilized mRNA was reconstituted in RNase-free water to a stock concentration of 1 µg/µL. and stored in single-use aliquots at −80 °C to prevent degradation.

Microinjection was performed using a FemtoJet 4i microinjector (Eppendorf, Hamburg, Germany) equipped with an M-152 micromanipulator (Narishige, Tokyo, Japan). Glass capillaries (BF100-50-10; Sutter Instrument) were pulled into fine needles using a P-97 micropipette puller. Approximately 5-10 pL of mRNA solution was injected into the cytoplasm of MI oocytes or the animal pole of one-cell stage embryos. Control groups were injected with an equivalent volume of RNase-free water or scrambled mRNA. Following injection, oocytes or embryos were rinsed three times and cultured in IVM developmental medium at 38.5 °C in a humidified atmosphere of 5% CO₂. The expression of the encoded protein was verified 12–24 h post-injection by immunofluorescence. Only embryos displaying normal morphology were used for subsequent phenotypic observations and time-lapse imaging.

### Sperm-Egg Binding and Polyspermy Analysis

Evaluation of Sperm Association with Zona-Free Oocytes To assess sperm-egg interaction, cumulus cells were removed from mature oocytes by incubation in 0.1% hyaluronidase. Denuded oocytes were subsequently treated with 0.1% Pronase (Roche Diagnostics, Basel, Switzerland) in TL-HEPES to digest the zona pellucida. Zona digestion was monitored under a stereomicroscope and terminated immediately upon dissolution of the zona. Zona-free oocytes were washed three times in TL-HEPES and subjected to IVF as described above. At 4 h post-insemination, oocytes were collected and rinsed three times in DPBS to remove non-specifically bound spermatozoa. Oocytes were stained with 10 µg/mL 4’,6-diamidino-2-phenylindole (DAPI), mounted on glass slides, and gently compressed under a coverslip. The number of spermatozoa adhering to the oocyte (plasma membrane) and those that had penetrated the ooplasm were quantified using fluorescence microscopy.

Assessment of Fertilization and Pronuclear Formation To evaluate the developmental progression and polyspermy rates, zona-free oocytes were prepared and fertilized as described above. At IVF-12 h, the resulting zygotes were collected, washed in DPBS, and stained with DAPI. Samples were mounted and examined under a fluorescence microscope. Fertilization was confirmed by the presence of at least one decondensed sperm head or pronucleus. The number of pronuclei (PN) per zygote was recorded to determine the rates of monospermic and polyspermic fertilization.

### Immunofluorescence

The following primary antibodies were used: Rat monoclonal anti-Folate Receptor 4 (JUNO; 1:50, Cat#ab228451; Abcam), rabbit polyclonal anti-GPI-PLD (GPLD1; 1:100, Cat#ab210753; Abcam). Secondary antibodies included Goat Anti-Rat, Alexa Fluor 594 (1:500,Cat#ab150160, Abcam, RRID:AB_2756445), and Donkey anti-Rabbit Highly Cross-Adsurbed Secondary Antibody, Alexa Fluor Plus 647 (1:500,Cat#A32795, Thermo Fisher Scientific, AB_2762835) and Donkey anti-Rabbit Highly Cross-Adsorbed Secondary Antibody, Alexa Fluor Plus 488 (1:500,Cat#A32790, Thermo Fisher Scientific, AB_2762833). Nuclei were counterstained with VECTASHIELD Antifade Mounting Medium with DAPI (1:500, Cat#H-1200, Vector, AB_2336790).

Oocytes were fixed in 4% paraformaldehyde (PFA) in PBS for 30 min and permeabilized with 0.5% Triton X-100 for 30–60 min at room temperature. Following blocking in 3% BSA in PBS for 1 h, oocytes were incubated with primary antibodies overnight at 4 °C. After three washes in PBS containing 0.1% Tween-20 (PBST), oocytes were incubated with appropriate fluorophore-conjugated secondary antibodies for 1 h at room temperature. For pronucleus and general morphology visualization, oocytes were washed in DPBS supplemented with 0.1% polyvinyl alcohol (PVA). Samples were mounted on non-fluorescent glass slides and imaged using a Nikon A2 laser scanning confocal microscope (Nikon, Tokyo, Japan).

### Image quantification and fluorescence intensity analysis

To ensure comparability, all images for a specific marker were acquired using identical confocal settings (laser power, gain, and offset). Quantitative analysis was performed using FIJI software (ImageJ; NIH, Bethesda, MD, USA). For each oocyte, a region of interest (ROI) was defined, and the mean fluorescence intensity was measured. Background subtraction was performed by measuring the signal in a non-stained cytoplasmic area or a negative control group. Data are expressed as average fluorescence intensity per unit area.

### Transmission Electron Microscopy (TEM)

For ultrastructural localization of JUNO and GPLD1, oocytes were fixed in a mixture of 4% paraformaldehyde and 0.2% glutaraldehyde in PBS for 1 h at room temperature. After three washes in PBS, free aldehydes were quenched with PBS/glycine, followed by blocking in 5% fetal calf serum (FCS) for 30 min. Oocytes were then incubated with primary antibodies (rat anti-JUNO or rabbit anti-GPLD1) for 1 h. After rinsing, specimens were incubated with a goat anti-rat secondary antibody conjugated to ultra-small gold particles (0.8 nm) for 30 min. Following secondary incubation, oocytes were washed and fixed in 1.5% glutaraldehyde for 30 min. Gold signals were subsequently amplified using a silver enhancement kit (Amersham IntenSE M; GE Healthcare) for 10 min, followed by rinsing in distilled water.

The specimens were post-fixed in 1% osmium tetroxide for 30 min, dehydrated through a graded ethanol series, and embedded in TAAB resin. Ultrathin sections (60 nm) were cut using a Leica UC6 ultramicrotome (Leica Microsystems, Wetzlar, Germany). Sections were collected on copper grids and contrast-stained with uranyl acetate and lead citrate. Grids were examined using a 120 kV FEI Tecnai Spirit BioTwin transmission electron microscope (FEI Company, Hillsboro, OR, USA) equipped with a HITACHI digital camera. Digital images were acquired to analyze the subcellular distribution of gold-labeled proteins.

### Live-cell time-lapse imaging

To dynamic visualization of protein localization, JUNO-eCFP mRNA was microinjected into the cytoplasm of MⅡ oocytes. Following injection, oocytes were washed extensively and cultured in 500 µL of maturation medium (IVM) in glass-bottom dishes for 12 h to allow for protein expression and completion of maturation.

To assess the effect of exogenous GPLD1, JUNO-eCFP-expressing oocytes were transferred to glass-bottom dishes containing TL-HEPES (Control) or TL-HEPES supplemented with human recombinant GPLD1 (Experimental). Imaging was performed using a laser scanning confocal microscope (Nikon A2) equipped with a stage-top environmental chamber maintaining 38.5 °C, 5% CO₂, and 100% humidity. Time-lapse images were acquired at a temporal resolution of 2 min for a total duration of 30min. Ten oocytes were recorded in parallel for each experimental session.

Spatiotemporal imaging of JUNO and GPLD1 during IVF. To track the coordination between JUNO and GPLD1 during fertilization, oocytes co-expressing JUNO-eCFP and GPLD1-eYFP were inseminated directly on the microscope stage. Processed boar spermatozoa were resuspended in mTBM and added to the fertilization droplets at a final concentration of 50000 sperm/mL (achieving a 500:1 sperm-to-oocyte ratio). For the inhibition assay, 1,10-phenanthroline (PHEN) was added to the mTBM droplets in the experimental group. Imaging commenced immediately upon insemination under stabilized environmental conditions (38.5 °C, 5% CO₂). Fluorescence intensities and protein recruitment patterns were analyzed from parallel recordings of 10–30 zygotes per group.

### Statistics and data analysis

Data were obtained from at least three independent biological replicates for each experimental condition. Statistical analyses were performed using GraphPad Prism 9 (GraphPad Software, San Diego, CA, USA). Differences between two groups were analyzed using an unpaired Student’s t-test, while multiple-group comparisons were conducted using one-way analysis of variance followed by Tukey’s or Sidak’s post-hoc test for multiple comparisons. All data are presented as the mean ±SEM. Statistical significance was defined as P < 0.05 (*), with P < 0.01 (**), P < 0.001 (***), and P < 0.0001 (****) representing increasing levels of significance.

### Online supplemental material

**Fig. S1** shows comparative analysis of protein expression and sequence conservation of JUNO and GPLD1 across mammalian species. **Fig. S2** shows concentration-dependent effects of PHEN on sperm–oocyte binding. **Fig. S3** shows construction and structural features of recombinant JUNO-eCFP and GPLD1-eYFP plasmids. **Fig. S4** shows fertilization-induced redistribution of JUNO and GPLD1. **Video 1** shows stable membrane localization of JUNO-eCFP in control group. **Video 2** shows rapid JUNO-eCFP shedding induced by recombinant hrGPLD1 protein. **Video 3** shows dynamic coordination of JUNO and GPLD1 during normal fertilization. **Video 4** shows inhibition of fertilization-induced JUNO shedding by PHEN. Provided online are two tables. **Table S2** shows significantly altered proteins in pairwise comparisons among pre- and post-fertilization stages, related to Fig. 4. **Table S3** shows a summary of upregulated and downregulated proteins across the three stages, related to Fig. 4.

## Supporting information

Figure S1. Comparative Analysis of Protein Expression and Sequence Conservation of JUNO and GPLD1 Across Mammalian Species.

Figure S2. Concentration-dependent effects of PHEN on sperm-oocyte binding.

Figure S3. Construction and structural features of recombinant plasmids.

Figure S4. Fertilization-induced redistribution of JUNO and GPLD1.

## Data availability

Data are available in the article itself and its supplementary materials.

## Acknowledgments

This research was funded by the National Natural Science Foundation of China (32372880), the Innovative Project of State Key Laboratory of Animal Biotech Breeding (Grant No. 2024SKLAB 1-6) and the 2115 Talent Development Program of China Agricultural University. The authors declare no competing financial interests.

## Author contributions

LZ and BDC conceived the study. LTS, FX, STG, and PYJ collected the oocyte and zygote samples. BDC and XGC analyzed the data with the help from XQZ and JYW. BDC and LZ drafted the manuscript, and LZ revised the manuscript with advice from GSL. LZ supervised the study and all authors read and approved the final manuscript.

---

Figure S1. **Comparative Analysis of Protein Expression and Sequence Conservation of JUNO and GPLD1 Across Mammalian Species. (A)** Relative protein expression levels of key sperm–oocyte recognition factors were compared among MII oocytes, IVF-4h, and IVF-24h stages. Several proteins exhibited stage-specific alterations, indicating dynamic regulation during fertilization. Data are presented as mean ± SEM. Statistical significance was determined by one-way ANOVA with multiple comparisons. *P < 0.05; **P < 0.01, ns, not significant. **(B)** Multiple sequence alignment of JUNO across five mammalian species. Sequences are aligned for mouse (Mus musculus), human (Homo sapiens), pig (Sus scrofa), sheep (Ovis aries), and cattle (Bos taurus). Identical residues are shaded in dark purple, while partially conserved residues are shaded in light purple. Green boxes indicate two predicted N-glycosylation sites, and the blue box denotes the C-terminal GPI-anchor signal sequence. **(C)** Amino acid sequence identity matrix for JUNO. The heatmap displays the pairwise sequence identity percentages among the five species. Color intensity correlates with the degree of similarity, with dark blue representing higher identity and light blue representing lower identity. **(D)** Multiple sequence alignment of GPLD1. Amino acid sequences from mouse, human, pig, sheep, and bovine are shown. Consistency in shading follows the convention in (B), with dark and light purple representing identical and partially conserved residues, respectively. **(E)** Amino acid sequence identity matrix for GPLD1. Pairwise identity percentages between the five species are shown. Values were determined via pairwise sequence alignment, where darker blue signifies higher sequence conservation.

Figure S2. **Concentration-dependent effects of PHEN on sperm–oocyte binding. (A)** Representative fluorescence images showing sperm bound to oocytes following treatment with increasing concentrations of PHEN (Control, 1, 10, 100, 250, 500, 750, 1000, and 1250 μM). Bound sperm are visualized by nuclear staining (blue). Scale bars = 100 μm. **(B)** Quantification of the number of sperm bound per oocyte across different PHEN concentrations. Data are presented as mean ± SEM. Statistical analysis was performed using one-way ANOVA followed by multiple comparisons. Asterisks indicate significant differences. *P < 0.05, ****P < 0.0001.

Figure S3. **Construction and structural features of recombinant plasmids.** Circular plasmid maps illustrating the organization of recombinant constructs JUNO-eCFP**(A)** and GPLD1-eYFP**(B)**, including promoter regions, gene inserts, selectable markers, and epitope tags. The direction of transcription is indicated by arrows. Relevant restriction enzyme sites and nucleotide positions are labeled for cloning validation.

Figure S4. **Fertilization-induced redistribution of JUNO and GPLD1. (A)** Representative confocal images showing the subcellular localization of JUNO-eCFP (cyan) and GPLD1-eYFP (yellow) in control group and PHEN-treated porcine oocytes at 0 h (MII stage) and 4 h post-IVF. **(B)** Quantitative analysis of JUNO-eCFP and GPLD1-eYFP fluorescence intensity in PHEN-treated zygotes.

Video 1. **Stable membrane localization of JUNO-eCFP in oocytes from control group.** Time-lapse imaging of porcine oocytes microinjected with JUNO-eCFP mRNA. The video shows consistent and stable fluorescence at the oocyte over a 30-minute observation period in the absence of exogenous GPLD1. Frames were captured every 45 seconds.

Video 2. **Rapid JUNO-eCFP shedding induced by recombinant hGPLD1 protein.** Time-lapse imaging of JUNO-eCFP-expressing oocytes following the addition of human recombinant GPLD1 (hrGPLD1) protein. The video demonstrates the progressive depletion and shedding of the JUNO-eCFP signal from the oocyte within 30 minutes, confirming the enzymatic cleavage of the GPI-anchor by GPLD1.

Video 3. **Dynamic coordination of JUNO and GPLD1 during normal fertilization.** Dual-color time-lapse imaging of oocytes co-injected with JUNO-eCFP (cyan) and GPLD1-eYFP (yellow) during in vitro fertilization (IVF) in The 240-minute recording.

Video 4. **Inhibition of fertilization-induced JUNO shedding by PHEN.** Time-lapse imaging of JUNO-eCFP and GPLD1-eYFP dynamics during IVF in the presence of PHEN in the 240-minute video.

