## Supplementary figures and images for "GPLD1 Regulates the Shedding of JUNO to Block Polyspermy in Porcine Oocyte"

### Figure S1. Comparative Analysis of Protein Expression and Sequence Conservation of JUNO and GPLD1 Across Mammalian Species.

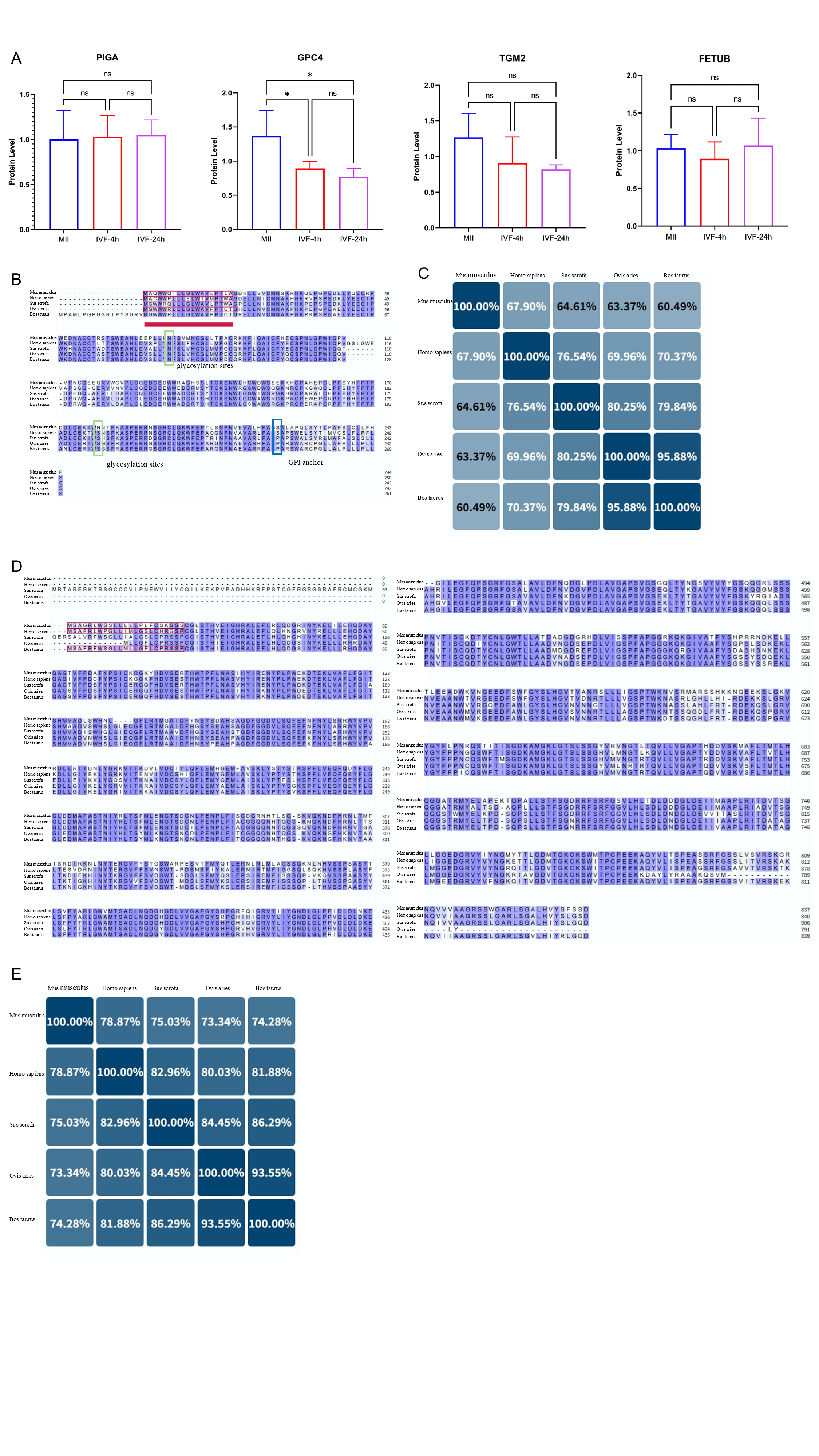

### Figure S2. Concentration-dependent effects of PHEN on sperm-oocyte binding.

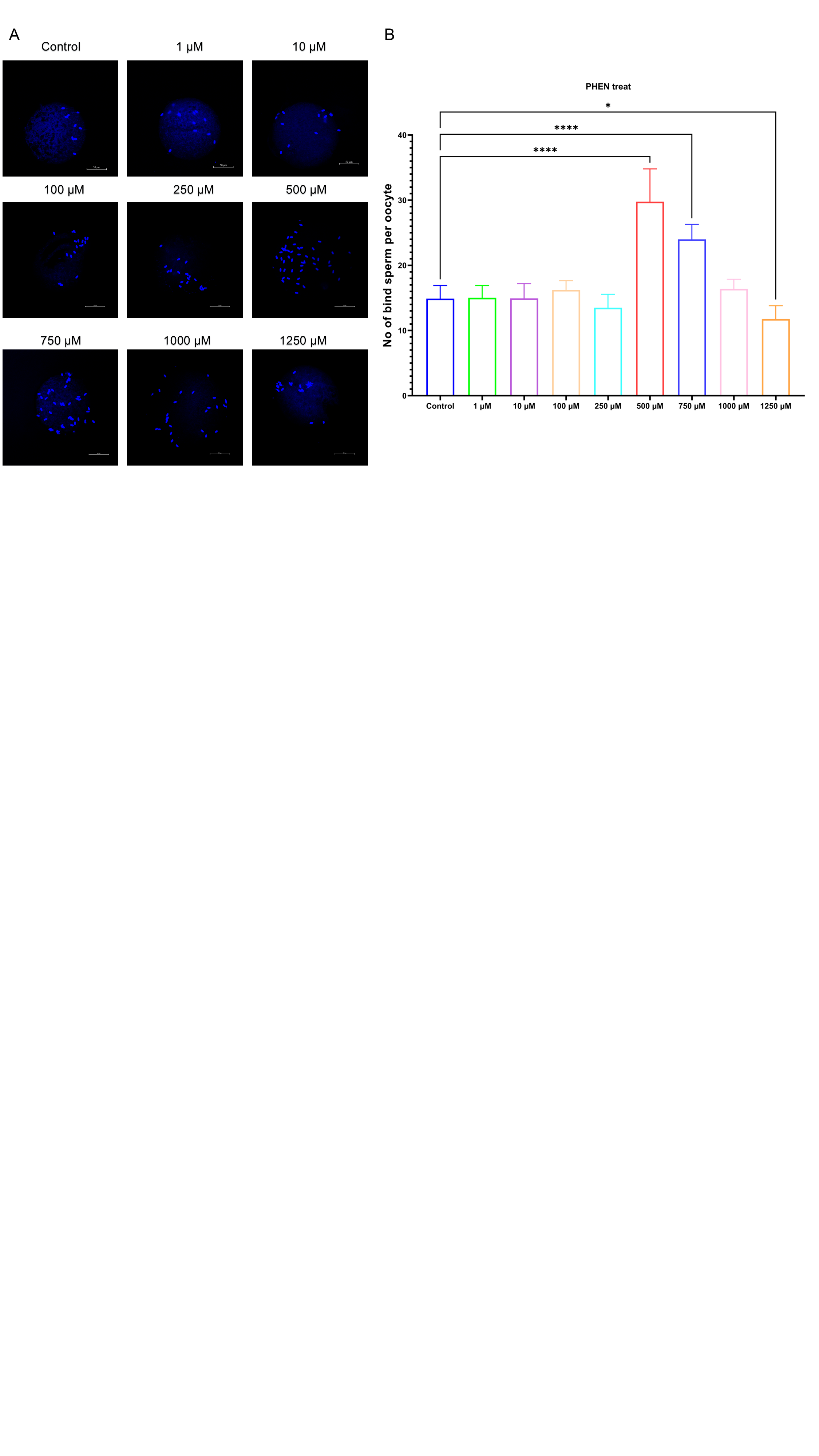

### Figure S3. Construction and structural features of recombinant plasmids.

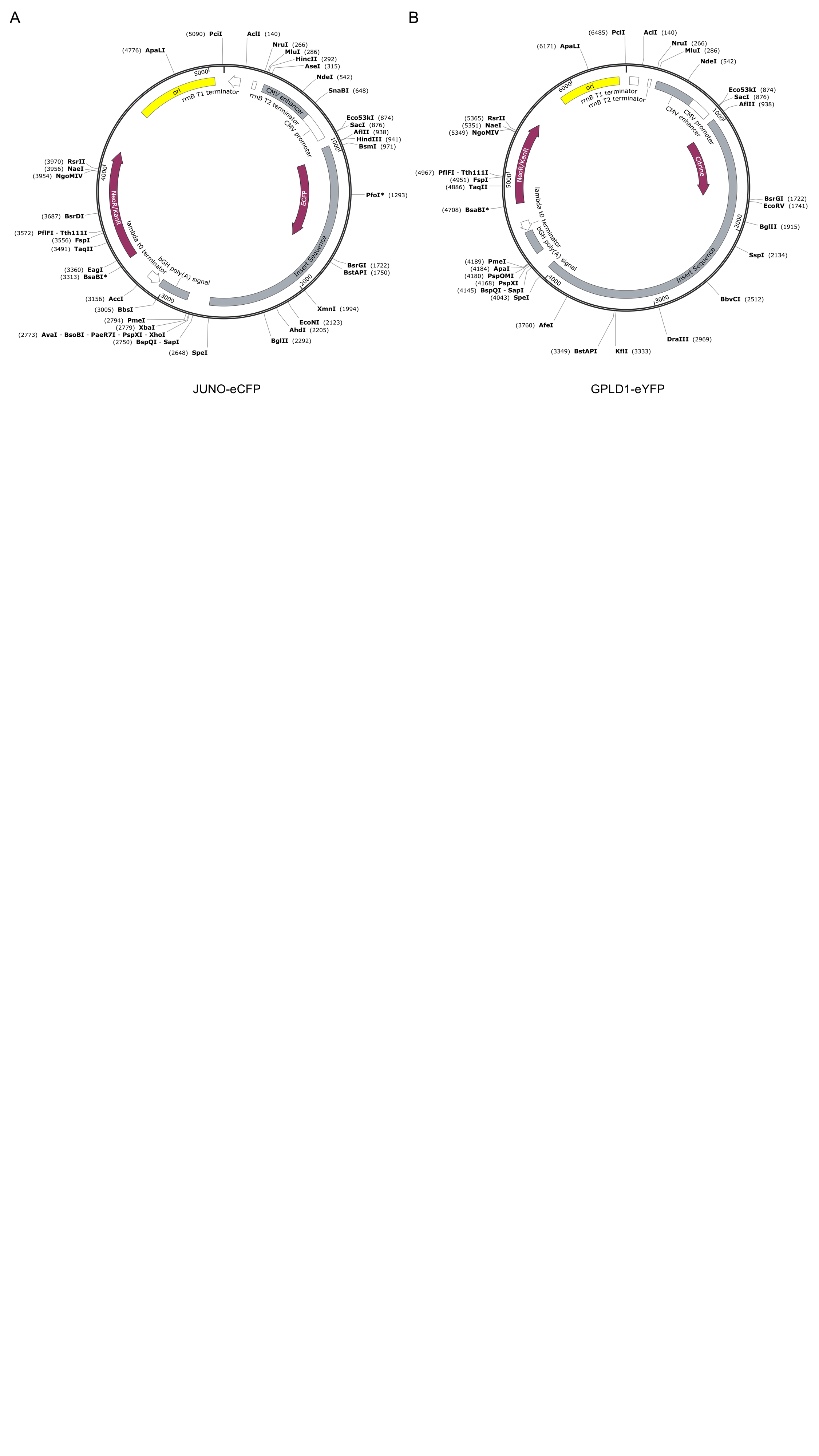

### Figure S4. Fertilization-induced redistribution of JUNO and GPLD1.

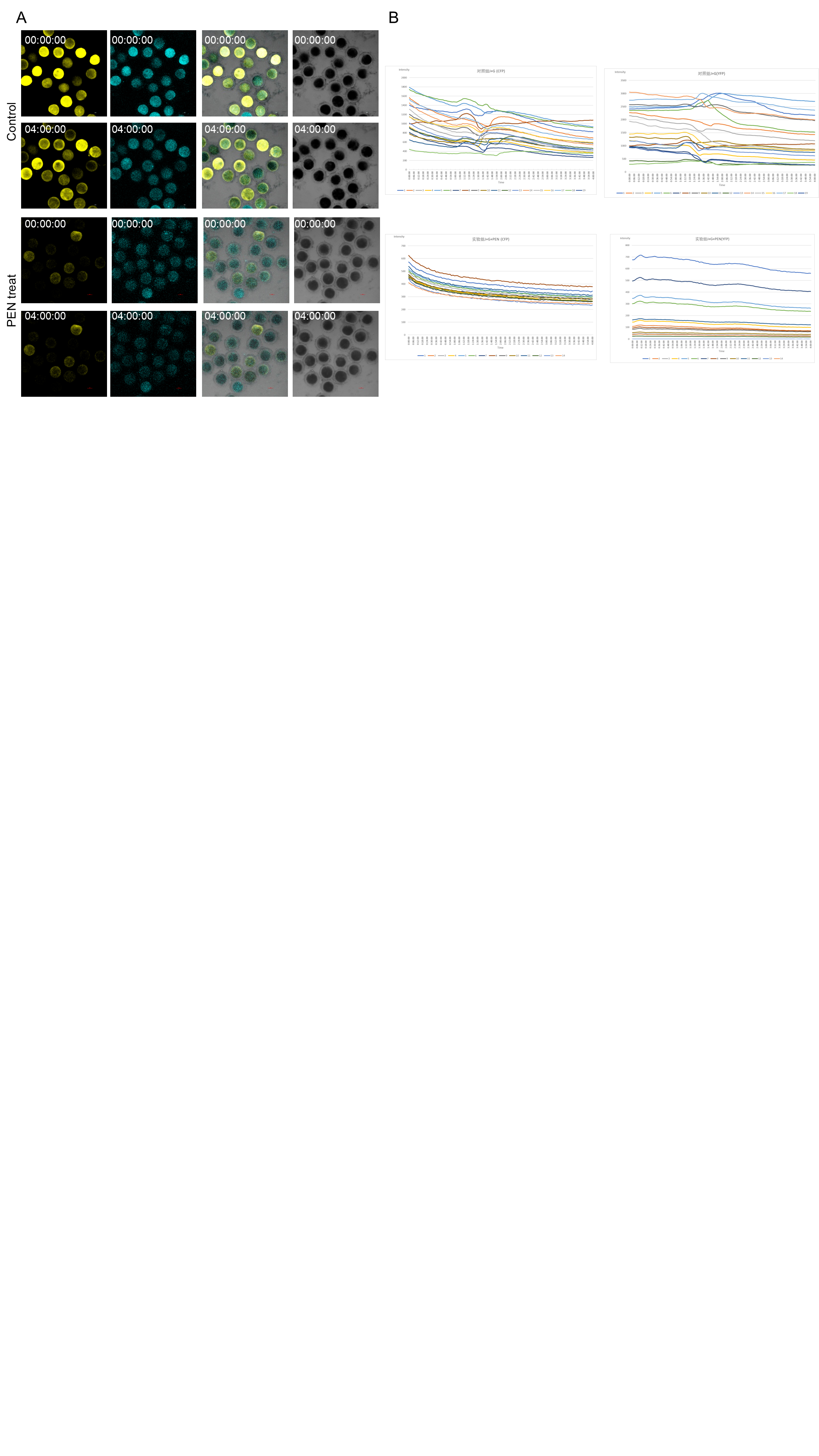
